# MRG proteins are shared by multiple protein complexes with distinct functions

**DOI:** 10.1101/2021.08.11.456009

**Authors:** Maëva Devoucoux, Céline Roques, Catherine Lachance, Anahita Lashgari, Charles Joly-Beauparlant, Karine Jacquet, Nader Alerasool, Alexandre Prudente, Mikko Taipale, Arnaud Droit, Jean-Philippe Lambert, Samer M.I. Hussein, Jacques Côté

## Abstract

MRG15/MORF4L1 is a highly conserved protein in eukaryotes that contains a chromodomain recognizing H3K36me3 in chromatin. Intriguingly, it has been reported in the literature to interact with several different factors involved in chromatin modifications, gene regulation, alternative mRNA splicing and DNA repair by homologous recombination. In order to get a complete and reliable picture of associations in physiological conditions, we used genome editing and tandem affinity purification to analyze the stable native interactome of human MRG15, its paralog MRGX/MORF4L2 that lacks the chromodomain, and MRGBP (MRG-binding protein) in isogenic K562 cells. We found stable interchangeable association of MRG15 and MRGX with the NuA4/TIP60 histone acetyltransferase/chromatin remodeler, Sin3B histone deacetylase/demethylase, ASH1L histone methyltransferase and PALB2/BRCA2 DNA repair protein complexes. These associations were further confirmed and analyzed by CRISPR-tagging of endogenous proteins and comparison of expressed isoforms. Importantly, based on structural information, point mutations could be introduced that can specifically disrupt MRG15 association with some complexes but not others. Most interestingly, we also identified a new abundant native complex formed by MRG15/X-MRGBP-BRD8-EP400NL that is functionally similar to the yeast TINTIN (Trimer Independent of NuA4 for Transcription Interactions with Nucleosomes) complex. Our results show that EP400NL, being homologous to the N-terminal region of NuA4/TIP60 subunit EP400, creates TINTIN by competing for BRD8 association. Functional genomics indicate that human TINTIN plays a role in transcription of specific genes. This is most likely linked to the H4ac-binding bromodomain of BRD8 along the H3K36me3-binding chromodomain of MRG15 on the coding region of transcribed genes. Taken together, our data provide a complete detailed picture of human MRG proteins-associated protein complexes which is essential to understand and correlate their diverse biological functions in chromatin-based nuclear processes.

**Highlights:**

- MRG15 and MRGX are stably associated with several different protein complexes important for genome expression and stability.
- Several MRG-containing complexes are chromatin modifiers.
- Specific point mutations in the MRG domain differentially affect associated complexes.
- A major human complex homologous to the yeast TINTIN complex is identified.
- The protein EP400NL competes with EP400 to functionally separate TINTIN from the NuA4/TIP60 complex.
- TINTIN contains a bromodomain and a chromodomain to regulate transcription.

## INTRODUCTION

The eukaryote genome is organized into a structure called chromatin, with the nucleosomes being its basic units, playing central roles in the regulation of all DNA-based processes for genome expression and maintenance (1). One major element governing chromatin structure and functions is the diverse post-translational modifications (PTMs) deposited on the external tails of the nucleosomal histones (2). PTMs on specific residues of specific histones include acetylation, methylation, phosphorylation and ubiquitination, which are recognized by dedicated reader domains in factors to regulate biological processes (3, 4). Disruption of the enzymes that “write” or “erase” these histone marks, or their “readers”, occurs in several human diseases, including cancer, leading to intense research efforts during the past several years to understand these regulatory interactions during cellular life (5).

One mark that has been studied extensively is the methylation of lysine 36 on histone H3 (H3K36me), which can occur as mono- (me1), di- (me2), or tri-(me3) methylation, levels being linked to distinct functions. While all methylation levels are generated by Set2 in *Saccharomyces cerevisiae* (6), mammalian methyltransferases like ASH1L and NSD1/2/3 only do H3K36me1/me2. Mammalian SETD2 on the other hand is responsible for H3K36me3 (7). Set2 associates with the elongating RNA polymerase II (RNAPII) phosphorylated on Ser2 of its C-terminal repeats (CTD). This leads to enrichment of H3K36me3 on the coding region of transcribed genes which recruits the RPD3S histone deacetylase complex to stabilize chromatin in the wake of RNAPII in order to avoid cryptic initiation and spurious transcription (8, 9). Mammalian SETD2 does not seem to affect global chromatin acetylation levels (7), even if H3K36me3 plays a central role in transcriptional activation and is distributed throughout the coding regions of active genes (10, 11). More precisely, H3K36me3 has been observed to be specifically enriched on exons, compared to introns, independent of nucleosome occupancy, in both yeast and mammals (12–14). These data were used to define a function of this histone mark in alternative mRNA splicing (15). It has been proposed that H3K36me3 is used as a platform recognized by the factor MORF-related gene on chromosome 15 (MRG15, a.k.a. MORF4L1) through its N-terminal chromodomain (CHD), an interaction that recruits the splicing machinery (16, 17). The yeast homolog of MRG15, Eaf3, is also known to affect splicing through its binding with the splicing factor Prp45/SKIP, thereby leading to the recruitment of spliceosome on intron-containing genes (18).

Mammalian MRG15 has a paralog expressed from the X-chromosome, MRGX (a.k.a. MORF4L2), which lacks the chromodomain. MRG15/X seem to regulate cell proliferation and senescence in specific contexts, while only MRG15 is required for mouse development (19, 20). They have been found associated with both lysine acetyltransferase (KAT) and deacetylase (KDAC) complexes, namely NuA4/TIP60 and Sin3B, also conserved in yeast (NuA4 and RPD3S) and containing the homologous protein Eaf3 (21–25). While MRGX does not possess a N-terminal CHD, it shares with MRG15 a highly conserved C-terminal MRG domain, crucial for their non-histone protein-protein interactions (20). Interestingly, it has been suggested that MRG15 can form an homodimer which is dissociated by its acetylation, regulating MRG15 association with other factors (26). MRG15 is also involved in DNA repair through its interaction with DNA repair factor PALB2, thereby favoring repair of double strand breaks by homologous recombination (27–29). It has been recently described that the MRG15-PALB2 complex is associated with undamaged chromatin on active genes to protect them from genotoxic/replicative stress (30). Finally, MRG15 was very recently implicated in diurnal rhythm of epigenomic remodeling through chromatin acetylation as well as lipid metabolism through gene activation with LRH-1 (31)

Although current knowledge on MRG15 points towards quite distinct important roles in the cell and during development, its described functions and associated cofactors are often studied in very artificial non-physiological conditions. Here, we aimed at defining the complete detailed interactomes of MRG15/X in the most native conditions possible. Tandem-affinity purification from isogenic cell lines identified several stable complexes containing MRG15 or MRGX. Endogenous protein tagging by CRISPR allowed validation and characterization of a new tetrameric protein complex with striking resemblance to the yeast TINTIN (<u>T</u>rimer Independent of <u>N</u>uA4 for <u>T</u>ranscription Interactions with <u>N</u>ucleosomes) complex that we have previously identified (25). The presence of a bromodomain and a chromodomain in this complex and its effect on transcription support the claim that we have identified a human TINTIN complex that associates with the body of active genes.

## EXPERIMENTAL PROCEDURES

### Recombinant Proteins and Pull-down Assay

Recombinant proteins (all human) 6xHis-BRD8, GST-MRG15 CHD long and short, GST-EP400(aa194-446), GST-EP400NL(aa51-297) and GST-ASH1L(aa2040-2634) were expressed in E. coli BL21 cells grown in LB media (from pET15b, pGEX4T3 or pGEX6P2 vectors). After induction with IPTG (30µM) overnight at 16°C, bacteria were harvested and GST or 6xHistagged proteins were purified as previously described with minor modifications (32).

Protein and chromatin pull-down experiments were performed as previously described (33). The GST pulldown assays were performed using 300 to 600 ng of GST-fused protein and an equivalent amount of His-tagged protein or 5μg of native H1-depleted chromatin from HeLa cells (prepared as described in (34)), which was precleared using glutathione-Sepharose beads. Equivalent protein levels were estimated through Coomassie blue-stained SDS-PAGE by comparison with known amounts of bovine serum albumin (BSA) standards. After pre-clearing, the His-tagged proteins were incubated with GST-immobilized protein in binding buffer (25 mM HEPES pH 7.9, 10% glycerol, 100 μg/ml BSA, 1 mM PMSF, 0.5 mM dithiothreitol (DTT), 0.1% Tween 20, and protease inhibitors, 250mM NaCl was used for chromatin pull-down) at 4°C for 1h to 4h followed by washing the beads three times. To visualize the proteins, the beads were loaded on SDS-PAGE gels, followed by Western blotting with anti-GST (Sigma G1160), anti-His (Clontech 631212), anti-H3K36me2 (Upstate 07-369), anti-H3K36me3(Abcam ab9050), anti-H3K79me3 (Abcam ab2621), and anti-H4 (Abcam ab7311). A GST-only protein and beads were used as a control.

### Experimental Model and Subject Details

K562 and U2OS cells were obtained from the ATCC and maintained at 37°C under 5% CO2. K562 were cultured in RPMI medium supplemented with 10% newborn calf serum and GlutaMAX. U2OS cells were cultured in DMEM medium supplemented with 10% fetal bovine serum.

### Establishment of Isogenic Cell Lines using the AAVS1 Safe Harbor or CRISPR/Cas9-mediated Genome Editing

K562 cells stably expressing 3xFLAG-2xStrep-tagged MRG15 spliced variants (long CHD or short CHD), MRGBP, MRGX, BRD8 spliced variants (one or two BRDs), EPC1 or EP400NL spliced variants (isoforms Q6ZTU2-5 and Q6ZTU2-6) were established as described before (35). BRD8, ASH1L, MRGX, and EP400NL were also endogenously 3xFlag-2xStrep-tagged using CRISPR/Cas9 with ouabain selection as previously described (36). The following gRNAs were used to generate these cell lines:

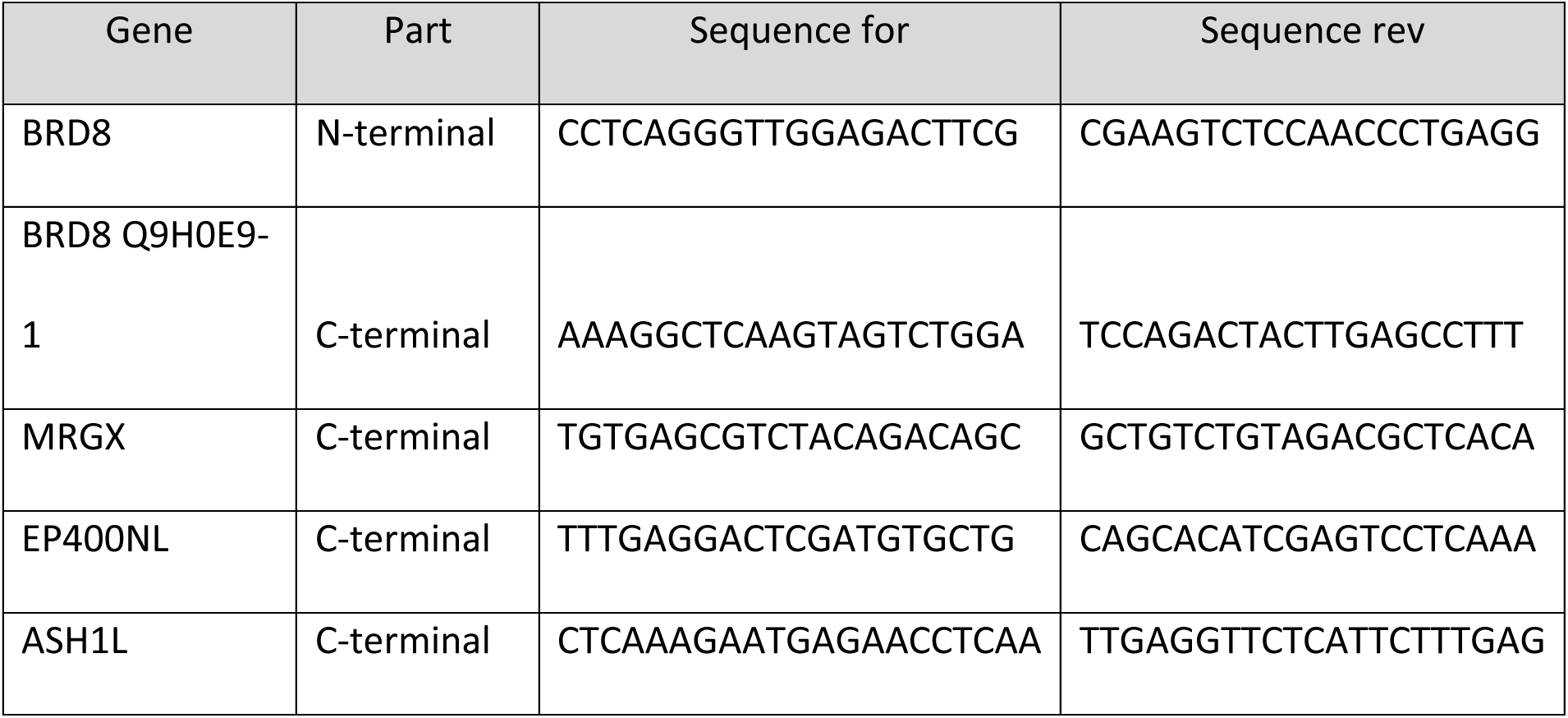

### Affinity Purification of Complexes

Native complexes were purified essentially as previously described (35). Briefly, Nuclear extracts were prepared from 2.5.10^9 cells and pre-cleared with CL6B sepharose beads. FLAG immunoprecipitations with anti-Flag agarose affinity resin (Sigma M2) were performed followed by two elutions with 3xFLAG peptide in elution buffer (20 mM HEPES-KOH [pH 7.9], 10% glycerol, 150 mM KCl, 0.1% Tween 20, 1mMDTT, 1 mM PMSF, 2 mg/mL Leupeptin, 5 mg/mL Aprotinin, 2 mg/mL Pepstatin, 10 mM Na-Butyrate, 10 mM β-glycerophosphate) with 200 μg/mL 3xFLAG peptide (Sigma). Then, STREP immunoprecipitations with Strep-tactin sepharose beads (Cedarlane) were performed followed by two elutions with elution buffer supplemented with 4mM biotin. Typically, 20µL of the first Strep elution was loaded on Nu-PAGE 4%–12% Bis-Tris gels (Invitrogen) and analyzed via silver staining. Fractions were then analyzed by mass spectrometry and western blotting. For purification after inducing DNA damage, cells were treated with 50ng/mL of neocarzinostatin (NCS) for 3h before beung collected to prepare the nuclear extract.

### Mass Spectrometry Analysis

To produce the peptides for mass spectrometry analysis, purified fractions were loaded on a 10% SDS-PAGE, ran until the dye migrated about 1 cm, stained with Sypro Ruby Red and a gel slice containing the entire protein signal was cut and processed for in-gel digestion with trypsin. Mass spectrometry analyses were performed by the Proteomics Platform of the CHU de Québec-Université Laval Research Center (Québec, Qc, Canada). Samples were analyzed by nano-LC/MSMS either with a 5600 triple TOF or an Orbitrap Fusion.

5600 triple TOF: Peptides were analyzed using an Ekspert NanoLC425 coupled to a 5600+ triple TOF mass spectrometer (Sciex, Framingham, MA, USA). Peptides were trapped at 4 μl/min in loading solvent (0.1% formic acid) on a 5mm x 300 μm C18 pepmap cartridge pre-column (Thermo Fisher Scientific / Dionex Softron GmbH, Germering, Germany) for 10 minutes. Then, the pre-column was switched online with a self-packed picofrit column (New Objective) packed with reprosil 3u, 120A C18, 15 cm x 0.075 mm internal diameter, (Dr. Maisch). Peptides were eluted with a linear gradient from 5-35% solvent B (A: 0.1% formic acid, B: acetonitrile, 0.1% formic acid) in 35 minutes, at 300 nL/min for a total run time of 60min. Mass spectra were acquired using a data-dependent acquisition mode using Analyst software version 1.7. Each full scan mass spectrum (400 to 1250 m/z) was followed by collision-induced dissociation of the twenty most intense ions. Dynamic exclusion was set for 12 sec and tolerance of 100 ppm.

Orbitrap fusion: Peptides were analyzed using a Dionex UltiMate 3000 nanoRSLC chromatography system (Thermo Fisher Scientific) connected to an Orbitrap Fusion mass spectrometer (Thermo Fisher Scientific, San Jose, CA, USA). Peptides were trapped at 20 μl/min in loading solvent (2% acetonitrile, 0.05% TFA) on a 5mm x 300 μm C18 pepmap cartridge pre-column (Thermo Fisher Scientific / Dionex Softron GmbH, Germering, Germany) for 5 minutes. Then, the pre-column was switched online with a Pepmap Acclaim column (ThermoFisher) 50 cm x 75µm internal diameter separation column and the peptides were eluted with a linear gradient from 5-40% solvent B (A: 0,1% formic acid, B: 80% acetonitrile, 0.1% formic acid) in 60 minutes, at 300 nL/min for a total runtime of 90 min. Mass spectra were acquired using a data-dependent acquisition mode using Thermo XCalibur software version 4.3.73.11. Full scan mass spectra (350 to 1800m/z) were acquired in the orbitrap using an AGC target of 4e5, a maximum injection time of 50 ms, and a resolution of 120 000. Internal calibration using lock mass on the m/z 445.12003 siloxane ion was used. Each MS scan was followed by MSMS fragmentation of the most intense ions for a total cycle time of 3 seconds (top speed mode). The selected ions were isolated using the quadrupole analyzer in a window of 1.6 m/z and fragmented by Higher energy Collision-induced Dissociation (HCD) with 35% of collision energy. The resulting fragments were detected by the linear ion trap at a rapid scan rate with an AGC target of 1e4 and a maximum injection time of 50ms. Dynamic exclusion of previously fragmented peptides was set for 20 sec and a tolerance of 10 ppm.

Database searching: MGF peak list files were created using Protein Pilot version 4.5 software (Sciex) for the data obtained with the 5600+ triple TOF and with Proteome Discoverer 2.3 software (Thermo) for the orbitrap data. The MGF sample files were then analyzed using Mascot (Matrix Science, London, UK; version 2.5.1). Mascot was set up to search a contaminant database and UniProtKB *Homo sapiens* database assuming the digestion enzyme trypsin. Mascot was searched with a fragment ion mass tolerance of 0.60 Da (orbitrap) or 0.1Da (5600+) and a parent ion tolerance of 10.0 ppm (orbitrap) and 0.1Da (5600+). Carbamidomethyl of cysteine was specified in Mascot as a fixed modification. Deamidation of asparagine and glutamine and oxidation of methionine were specified in Mascot as variable modifications. 2 missed cleavages were allowed.

Criteria for protein identification: Scaffold (version Scaffold_4.8.7, Proteome Software Inc., Portland, OR) was used to validate MS/MS-based peptide and protein identifications. Peptide identifications were accepted if they could be established at greater than 8.0% probability to achieve an FDR less than 1.0% by the Scaffold Local FDR algorithm. Protein identifications were accepted if they could be established at greater than 99.0% probability to achieve an FDR less than 1.0% and contained at least 2 identified peptides. Protein probabilities were assigned by the Protein Prophet algorithm (37). Proteins that contained similar peptides and could not be differentiated based on MS/MS analysis alone were grouped to satisfy the principles of parsimony. Data were compared to mock purified fractions obtained from cells expressing an empty TAP-tag from the *AAVS1* site. Data were further analyzed using the CRAPome online tool and filtered against similar experiments (www.crapome.org). For all samples analyzed, protein identifications, corresponding accession numbers, numbers of spectral counts/distinct peptides and percentage of protein coverage are listed in supplemental Excel tables included in one workbook file.

All MS files generated as part of this study were deposited at MassIVE (http://massive.ucsd.edu). The MassIVE ID is MSV000087245 and the MassIVE link for download is http://massive.ucsd.edu/ProteoSAFe/status.jsp?task=9534fb8b77634860938b1139a53d9df8. The password for download prior to final acceptance is MRG15.

### Antibodies and siRNAs

The following antibodies were used for Western blotting at the indicated dilution: anti-FLAG - HRP conjugate (Sigma M2, 1:10000); anti-Brd8 (Bethyl A300-219A, 1:10000); anti-DMAP1 (Thermoscientific PA1-886, 1:1000); anti-P400 (Abcam ab5201, 1:1000); anti-MRG15 (Active Motif 39361, 1:1000); anti-MRGBP (Abnova H00055257-B01, 1:2000); anti-MRGX (Abnova PAB6152, 1:1000); anti-KAT5/Tip60 (Abcam ab137518, 1:1000); anti-ASH1L (Bethyl A301-749, 1:1000); anti-KDM5A (Cell signaling 3876, 1:1000); anti-Pf1 (Bethyl A301-647, 1:1000); anti-PALB2 (Bethyl A301-246, 1:3000); anti-AKAP8 (Abcam ab72196, 1:2500); anti-GAPDH (Thermofisher 39-8600, 1:10000); anti-KAP1 (MAB3662, 1/4000).

The following siRNA were used against the indicated proteins: KAT5/TIP60 smartpool (Dharmacon, CCACAGAUCACCAUCAAUG, GAACAAGAGUUAUUCCAG, CACAGGAACUCACCACAUU, GGACAGCUCUGAUGGAAUA), BRD8 from Sigma (SASI_Hs01_00131635; target sequences start at nucleotide 2223); MRG15 smartpool (Dharmacon, GAAGAGCCUUGCUUUAUUA, UAAAGUACCUGGCAAAGAA, GUACCCAGCUACUCUAUAA, GAUGACUGGGACUUAAUUA); MRGX smartpool (Dharmacon, GCACUCAGCUGCUCUACAA, GAGGAGGCGUUUAAGAAUA, GCAGGGAAAUGUUGAUAAU, GCUCCAAUGUCCCAGGUUU), MRGBP smartpool (Dhaarmacon, GAAGAACUCCUCAGACUUG, GAACUUCGUCCUUCCAGAA, GAACCGACACUUCCACAUG, ACAAAGUCCUGACCGCAAA), and siLuciferase as control is used (DOYYA, AACUGACGCGGAAUACUUCGA). Knockdown efficiency was validated by RT-qPCR with the following primers:

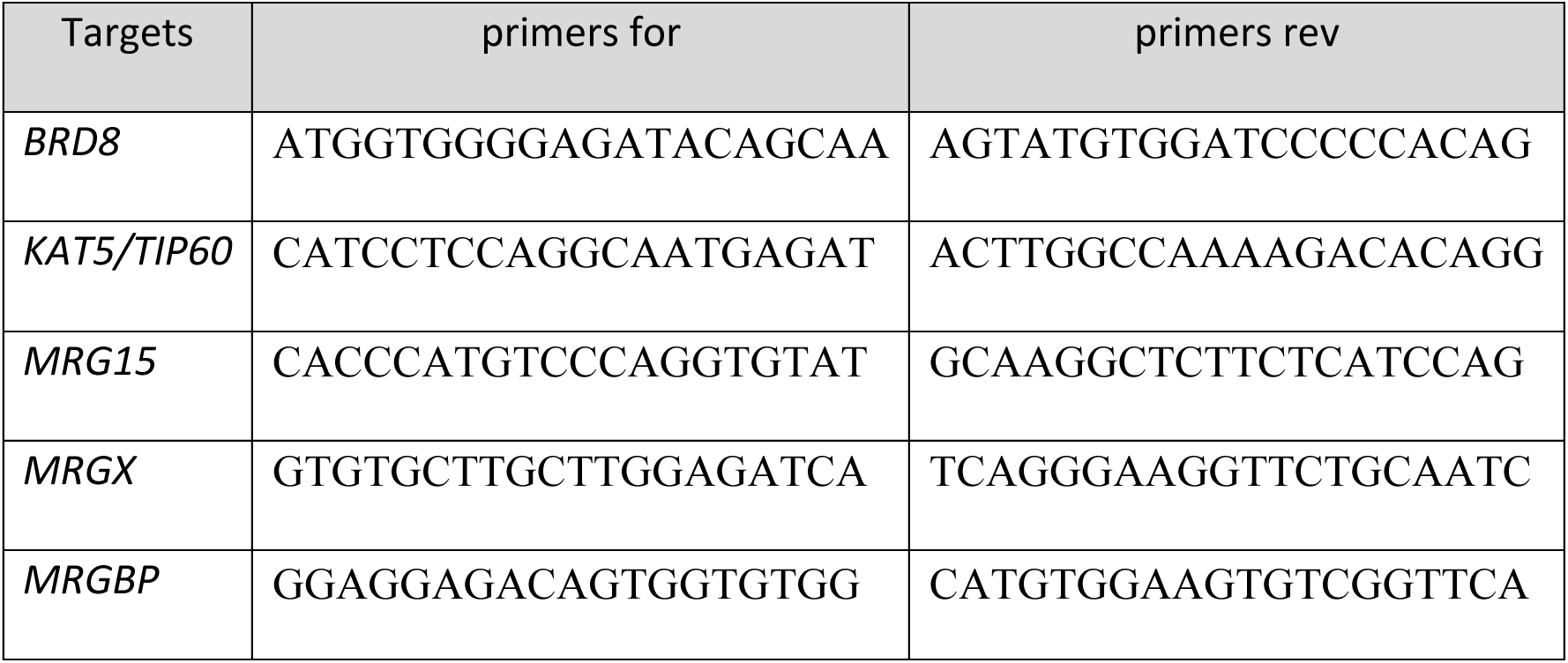

### In Vitro Histone Methyltransferase Assay

Purified endogenous ASH1L (or truncated recombinant GST-ASH1L(aa2040-2634)) was first preincubated with recombinant MRG15 purified from baculovirus (or the same amount of BSA) and 0.5µg of native Short OligoNucleosomes (SON) purified from HeLa cells for 4h at 30°C in a 20µl reaction containing 100mM Tris-HCl pH8.0, 25% glycerol, 0.5mM EDTA, 5mM DTT, 5mM PMSF, 10mM MgCl_2_ for 30min at 4C. Then, 2 µL of ^3^H S-adenosyl methionine (0.55uCi/ul) was added and the mixture was incubated for 3h at 30°C. the reaction was then spotted onto p81 filter paper, washed in carbonate buffer and scintillation counting was used to determine the incorporation.

### RNA-sequencing Analysis

2 days post-Knockdown (KD) of MRG proteins, BRD8 or KAT5 in U2OS cells, RNA was extracted with the Monarch Total RNA Miniprep Kit (NEB #T2010), following the manufacturer’s indications. Then, at least 1.5μg of RNA were depleted of rRNAs using the NEB rRNA-depletion kit (HMR) followed by Illumina library preparation. Finally, cDNAs were sequenced using the NovaSeq6000 S2 for 50M of PE100 reads. Two replicates of each KD were sequenced. Raw sequences and processed data of short reads RNA-sequencing from U2OS cells were deposited in the GEO database under accession number **GSE181533**.

### Gene expression analysis

Raw fast5 files were base called using Guppy v.4.2.2 with the guppy_basecaller command and the dna_r9.4.1_450bps_fast.cfg configuration file and default settings. Barcodes were detected and reads were separated with the guppy_barcoder using the default settings. The resulting FASTQ reads were aligned to the hg19 human reference with Minimap2 with the following parameters: -aLx splice --cs=long. Raw read counts were obtained with the featureCounts tool from the Subread package v 2.0.0, using the exon counting mode (38, 39). EdgeR R-package (v3.12.1) was then used to normalize the data, calculated RNA abundance at the gene and transcript levels (as counts per million reads (CPM), and perform statistical analysis (40). Briefly, a common biological coefficient of variation (BCV) and dispersion (variance) were estimated based on a negative binomial distribution model. This estimated dispersion value was incorporated into the final EdgeR analysis for differential gene expression, and the generalized linear model (GLM) likelihood ratio test was used for statistics, as described in EdgeR user guide. Analysis of variations in gene expression was performed by comparing KDs BRD8, MRGBP, MRG15, MRGX, and KAT5/TIP60 to siControl (siLuciferase). From all genes showing differential expression, a cutoff of 2-fold difference was applied to the Log2(fold change) values between the selected sample and control.

### mRNA splice variants analysis

Reads were trimmed using fastp v0.20.0 (41). Quality check was performed on raw and trimmed data to ensure the quality of the reads using FastQC v0.11.8 and MultiQC v1.8 (42, 43). The quantification was performed with Kallisto v0.46.2 (44). Differential expression analysis was performed in R v4.0.0 using the DESeq2 v1.28.1 (45, 46). Analysis of variations in splice isoform expression was performed by comparing differential expression at the gene and transcript levels. From all genes showing no differential expression with the selected control (padj ≥ 0.1), those with significant differences in transcript expression (padj ≤ 0.1) were selected. To highlight the genes with high changes in transcript expression compared to gene expression, a cutoff of 2-fold difference was applied to the Log2(fold change) values between the selected sample and control.

### Reverse Transcription-qPCR

siRNAs for KD of BRD8 or MRG15 or MRGBP or MRGX or KAT5/TIP60 were transfected in U2OS cells. siLuciferase (siLuc) was used as control. 48hrs later total RNA was extracted with the RNeasy Plus Mini kit (QIAGEN). 500 ng of RNA was reverse transcribed by oligo-dT and random priming into cDNA with a qScript cDNA SuperMix kit (QuantaBio-VWR), according to the manufacturer’s instructions. Quantification of the amount of cDNA was done with SYBR Green I (Roche) on a LightCycler 480 (Roche) for real-time PCR. The *RPLP0* gene was used as a housekeeping internal control for normalization. The error bars represent the range based on independent experiments. The oligonucleotide sequences used for expression analysis by RT-qPCR are listed below:

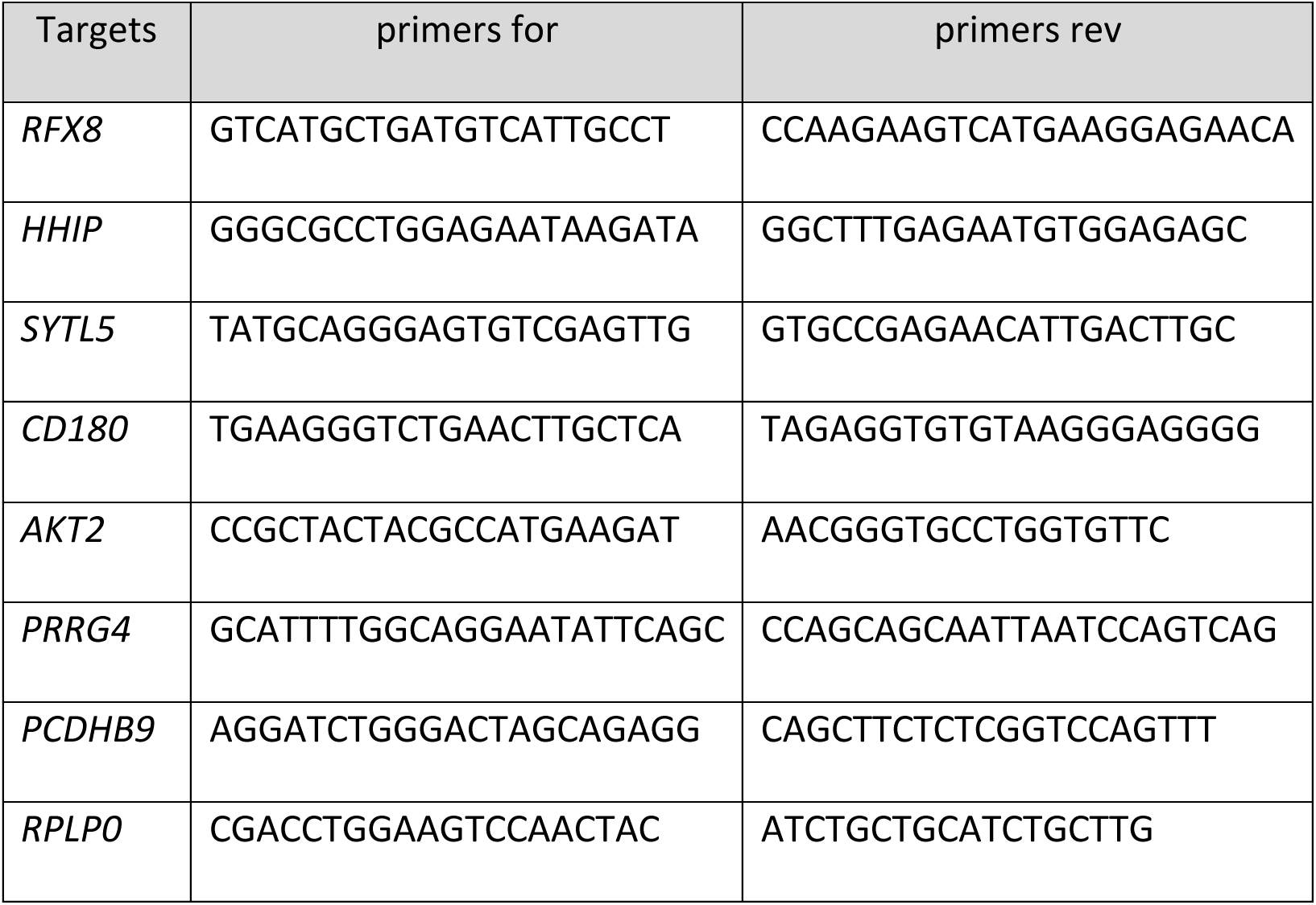

### Experimental Design and Statistical Rationale

A mock K562 cell line expressing an empty 3xFlag-2xStrep tag from the *AAVS1* locus is used as control. Most purification of tagged proteins shown are from single representative replicate experiments, while protein IDs (and loss in mutants) were carefully validated in purified fractions by gels and western analyses. Wild type MRG15S purified fractions included are from 3 biological replicates while the W172A/Y235A double mutant are from 2 biological replicates. Each Knockdown in U2OS cells was performed in biological duplicates. Control with a siLuciferase was generated concomitantly to experimental samples. RT-qPCRs to validate Knockdown and RNA-sequencing were performed in biological duplicates. Error bars represent the range from two biological replicates. Statistical analyses were performed via two-way ANOVA using Prism version 7 (GraphPad software inc California, USA) followed by Tukey’s test. P-values <0.05 were considered significant.

### Recruitment-activator assay

This assay is described in (47). The reporter HEK293T cell line with a TRE3G-EGFP reporter was a kind gift from Stanley Qi (Stanford University). The reporter cell line was transduced with a lentivirus expressing ABI-dCas9 followed by two rounds of selection under 6µg/ml blasticidin. These cells were then transduced with a lentivirus expressing EBFP2 and a gRNA targeting seven tetO repeats in the TRE3G promoter (gRNA sequence GTACGTTCTCTATCACTGATA). Single cell derived clonal cell lines were generated and a clone showing robust EGFP induction by a strong transcriptional activator VPR was selected for downstream assays. 96-well plates were seeded with 3x10^4^ cells per well one day prior to transfection. 150 ng of each construct was transfected using polyethylemine (PEI). Transfected cells were induced 24 hours after transfection by treatment with 100 µM abscisic acid. 48 hours after induction, cells were dissociated and resuspended in flow buffer using a liquid handing robot and analyzed by LSRFortessa (BD). Flow cytometry data was analyzed using FlowJo by gating for positive gRNA (EBFP2), then further for construct (TagRFP) expression. At least 25,000 cells were analyzed for each replicate.

## RESULTS

### Impact of the CHD on the binding of MRG proteins to chromatin

Although MRG15 is highly conserved from yeast to human, it is interesting to observe that two isoforms are produced in mammals through alternative mRNA splicing and that a paralogous protein, MRGX, is produced by another gene. Interestingly, the major difference between these 3 proteins resides in the chromodomain (CHD). MRG15 isoforms contain either a “long CHD” (Q9UBU8-1 in UniProtKB) with an extra 39 amino acids (aa) insert or a “short CHD” (Q9UBU8-2 in UniProtKB) which is considered canonical. In contrast, MRGX lacks the CHD region at its N-terminus (**Fig. 1A**). Yeast Eaf3 in comparison also has a “long CHD” but the extra insert is located on the opposite side of the hydrophobic cage that recognizes H3K36me3 compared to MRG15 (**Fig. S1A**). The proteins are characterized by a ∼175aa MRG domain at their C-terminus, important for their non-histone protein-protein interactions (20). Eaf3 CHD has been shown to bind H3K36me2/me3 which is required for suppression of spurious intragenic transcription by the RPD3S histone deacetylase complex (24,48,49). While Eaf3 CHD has been argued to also bind H3K4me3 *in vitro*, MRG15 short CHD does not show any interaction with this mark (50). We then wondered if the insert in MRG15 long CHD isoform could change its binding properties. To test that, we performed GST pull-down experiments with purified native chromatin and recombinant proteins containing the short or long CHD of MRG15. The short CHD showed great specificity for H3K36me3, compared to H3K36me2, while the long CHD seem to have weaker affinity (**Fig. 1B**).

**Figure 1.**
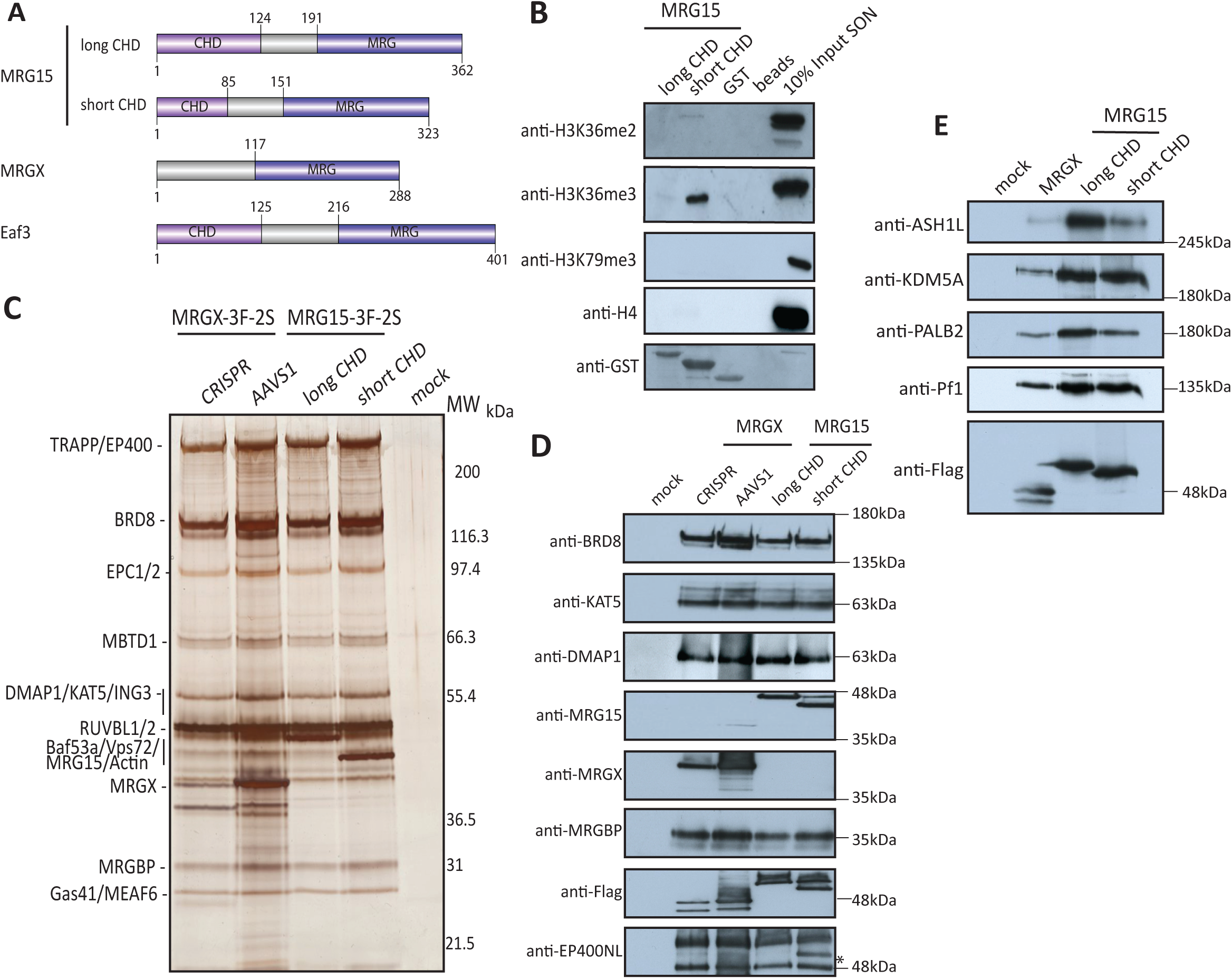
The interactome of MRG proteins highlights the association with several distinct complexes. (A) Schematic representation of MRG15 spliced variants, MRGX, and the yeast Eaf3 functional domains. (B) Short chromodomain MRG15 binds preferentially H3K36me3 *in vitro*. Recombinant GST-tagged MRG15 isoforms or GST or beads were incubated with LON followed by washes and western blotting. Anti-H4 and anti-H3K79me3 are used as control. (C) 3xFlag-2xStrep elutions of indicated purifications from K562 cells were migrated on 4-12% SDS-PAGE and analyzed by silver staining. MRGX and MRG15 spliced variants bind the acetyltransferase NuA4/TIP60 complex. Subunits were identified by mass spectrometry (Table S1). Mock is a purification from cells expressing an empty tag. (D-E) Purified complexes from C) were analyzed by western blot with the indicated antibodies to confirm the equivalent presence of known subunits of the NuA4/TIP60 complex (D) and the other known interactors of MRG proteins (E) (* non-specific band).

To further examine the potential impact of the CHD size and or its absence on chromatin binding *in vivo*, we generated isogenic K562 cell lines with a single copy of the epitope-tagged (3xFlag-2xStrep) MRG15 isoforms or MRGX cDNA inserted at the *AAVS1* safe-harbor locus, as described before (**Fig. S1B**)(35). We then analyzed their genome-wide locations using ChIP-seq experiments with an anti-Flag antibody. MRG15-S, MRG15-L and MRGX seem the share the vast majority (>70%) of their genomic targets, independently of the CHD status (**Fig. S1C**). On the other hand, MRG15-L had a relatively much smaller number of bound regions that could be mapped, possibly reflecting a looser interaction with chromatin. It remains to be seen if the small proportion of genomic regions uniquely bound by MRGX or MRG15 are functionally significant.

### MRG proteins are associated with several distinct protein complexes but are interchangeable

As mentioned above, several associated factors have been identified for mammalian canonical MRG15 (short CHD/MRG15-S), often using methods like transient or stable over-expression and simple immunoprecipitations. Thus, we wanted to get a full picture of the associated complexes in the most native conditions using our validated approaches. At the same time, we investigated if MRG15-L, MRG15-S and MRGX have different interactomes. Even though the *AAVS1* system usually allows near physiological level of expression (within 2-2.5-fold (35)), it does not recapitulate native regulation by endogenous promoters and 5’/3’UTRs. We were able to tag the endogenous *MRGX* gene using an improved CRISPR/Cas9 genome editing method (36). We used a gRNA targeting the stop codon of MRGX to introduce the 3xFlag-2xStrep tag (**Fig. S1D**). Comparing endogenous versus *AAVS1*-mediated expression levels indicates significantly lower endogenous expression, but similar to MRG15 expression from AAVS1 (**Fig. S1B**).

We then performed tandem affinity purifications with native elution at both steps, as previously described (35, 51). The final elutions with biotin were then analyzed by silver staining, mass spectrometry and western blotting (**Fig. 1C-E**, **Table S1**). In agreement with the highly conserved MRG domain between MRG15 and MRGX, we observed very similar interactomes for all MRG proteins. We could detect the full set of subunits for the NuA4/TIP60 acetyltransferase complex as well as the Sin3B deacetylase complex (homologous to yeast RPD3S) that includes the KDM5A H3K4 demethylase. It is interesting to point out that MRG15 and MRGX proteins seem largely mutually exclusive in their associations since we could detect only 1 or 2 spectral counts specific for MRGX in MRG15 preps and vice versa (**Fig. 1D** and **Table S1**). The data is also arguing that the reported ability of the MRG domain to homodimerize in an acetylation-regulated manner is not a major event (26). MRG15-S and MRGX have also been previously linked to PALB2/BRCA2, playing a crucial function in DNA repair by homologous recombination (27,29,30) and we confirmed that (**Fig. 1E**, **Table S1**). This interaction seems stable and not modulated by a cellular DNA damage response since we obtained very similar results purifying MRG15-S from cells treated with DNA double-strand break-inducing agent neocarzinostatin (NCS), for PALB2/BRCA2 or any other interacting partners (**Fig. S2A**, **Table S1**), in agreement with the proposed role of MRG15/PALB2/BRCA2 constitutive association in protecting undamaged chromatin on active genes from replication-associated stress (30). MRFAP1 is also an interesting interactor of both MRG15 spliced isoforms and MRGX as it was known to form a complex with MRG15-S which is regulated by NEDDylation (52). Finally, a particularly interesting shared interaction is with the H3K36 methyltransferase ASH1L (**Fig. 1E**, **Table S1**).

### Endogenous ASH1L is associated with MRG proteins

ASH1L is a member of the trithorax group proteins involved in activation of Hox genes in *Drosophila*. Its methyltransferase activity is crucial for H3K36me1/me2 maintenance, thereby leading to MLL recruitment on its specific promoter targets (53, 54). The catalytic activity of ASH1L seems also important for DNA repair via nucleotide excision (55). Moreover, previous studies have shown that MRG15 and MRGX enhance the methyltransferase activity of a truncated version of ASH1L (56, 57). Indeed, this binding can affect the auto-inhibitory loop of ASH1L (58, 59). Importantly, this stimulatory effect is therefore independent of the H3K36me-binding CHD of MRG15. To confirm this association and biochemical studies in the physiological context, we were able to tag and purify endogenous ASH1L using the CRISPR/Cas9 system in K562 cells (**Fig. 2A**, **S1E**). We then analyzed by western blots and mass spectrometry the ASH1L interactome. We confirmed association of MRG15 and MRGX as well as Nurf55/RBBP4, an already known factor link to ASH1L (**Fig. 2B**, **Table S2**). Interestingly, the A Kinase Anchoring protein 8 (AKAP8) and multiple RNA-processing factors were also found associated with ASH1L, suggesting a potential function in alternative splicing, as it was already described for AKAP8 (60).

**Figure 2.**
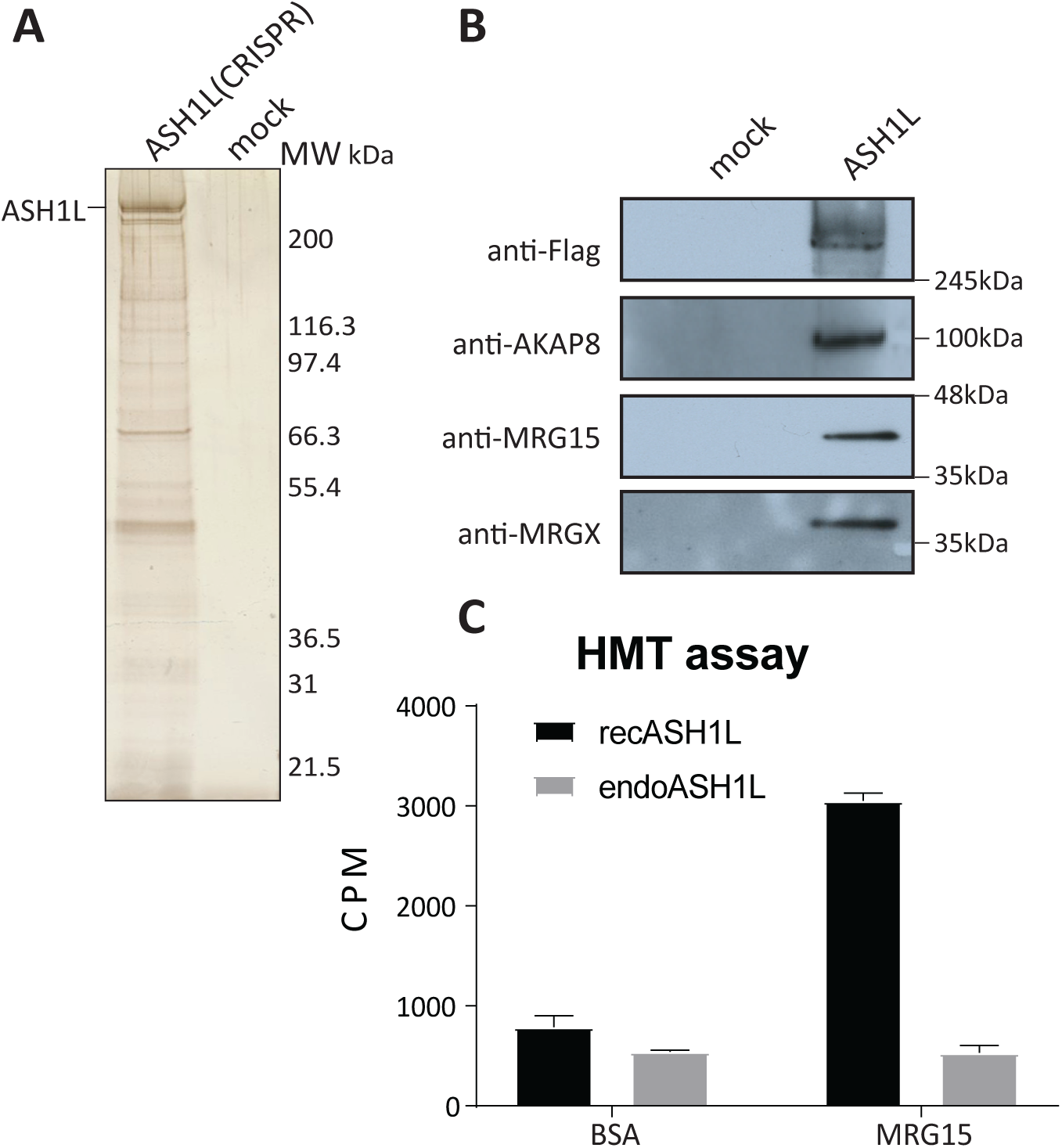
Endogenous ASH1L is stably associated with MRG proteins. (A) Purification of endogenous ASH1L from K562 cells modified by CRISPR/Cas9 system to add 3xFlag 2xStrep. The elution was analyzed on 4-12% SDS-PAGE followed by silver staining. (B) Purified ASH1L from A) was analyzed by western blot with the indicated antibodies to confirm the interaction with ASH1L and its interactors such as MRG proteins and AKAP8. (C) *In vitro* histone methyltransferase assay performed with recombinant ASH1L (aa2040-2634)- pGEX6P2 (recAHS1L) and strep elution from endogenous ASH1L (endoASH1L). The graph shows the scintillation counts of the liquid assays with 3H S-adenosyl methionine (SAM) with native short oligonucleosomes (SON). An empty vector pGEX6P2 and a mock cell line were used as control and subtracted from final counts. Error bars represent the range of two technical replicates.

To confirm the role of MRG15 in stimulating ASH1L methyltransferase activity, we performed an *in vitro* histone methyltransferase assay (HMT) to measure its activity. We used a recombinant truncated version of ASH1L (aa 2040-2684), preincubated with recombinant MRG15 or BSA. As expected, MRG15 greatly enhanced the methyltransferase activity of ASH1L (**Fig. 2C**). In parallel, the methyltransferase activity of the purified endogenous ASH1L was not affected by the preincubation with recombinant MRG15. This strongly suggests that the vast majority of endogenous ASH1L is stably bound to an MRG protein *in vivo,* being fully active and non-responsive to addition of exogenous MRG15.

### Genetic mutations to modulate MRG15 association with distinct complexes

Previous work has been done to characterize the structure of the MRG domain as an interface for protein-protein interactions (20,58,59,61,62). Based on this information, we investigated if we could manipulate the association of MRG proteins with distinct complexes in native conditions. We generated K562 cell lines expressing tagged MRG15 carrying the W172A and Y235A substitutions in its MRG domain, which have been shown to affect some interactions (PF1, MRGBP, ASH1L) (20)in binary recombinant experiments (**Fig. 3A**)(20,59,61). After purification and analysis by silver-stained gel, western blot and mass spectrometry, we can conclude that these mutations specifically disrupt MRG15 association with the Sin3B deacetylase complex, BRCA2/PALB2, MRFAP1 and ASH1L (**Fig. 3B-C**, **S2B**, **Table S3**). Strikingly, MRG15 association with the NuA4/TIP60 complex is still very stable based on western signals, maybe a slight decrease in recovery based on spectral counts. This clearly indicates that distinct molecular features of the MRG domain allow the multi-specificity of protein interactions and therefore associations to complexes. These mutants could help address the functional implication of MRG proteins association with these partners versus its presence in NuA4/TIP60. To see if it would be possible to address the opposite question, we again used published structural information (20) and introduce point mutations in the MRG-binding protein MRGBP, a subunit of NuA4/TIP60 directly associated with MRG15/X in the complex. We generated K562 cell lines expressing tagged wild-type and mutant MRGBP carrying the W78A F105A substitutions, mutations known to affect MRG15 binding in binary recombinant experiments (**Fig. 3A**)(20). Analysis of the purified fractions showed that the double mutant disrupted the association of MRG15 and MRGX with the native NuA4/TIP60 complex without affecting other components (**Fig. 3D-E**). Together, these MRG mutants have the potential to help us understand the specific function of MRG protein in distinct complexes. For example, as mentioned before, MRG15 is known to have a function in the repair of DNA double-strand breaks by homologous recombination, certainly in part through its association with PALB2/BRCA2 (27,29,30,63). Furthermore, the NuA4/TIP60 complex plays also plays a role in DNA repair, promoting homologous recombination (64–66). Thus, taking advantage of its mutants, it will be possible to characterize the function of MRG15 in DNA repair, distinguishing its role from within the NuA4/TIP60 complex versus its association with PALB2/BRCA2.

**Figure 3.**
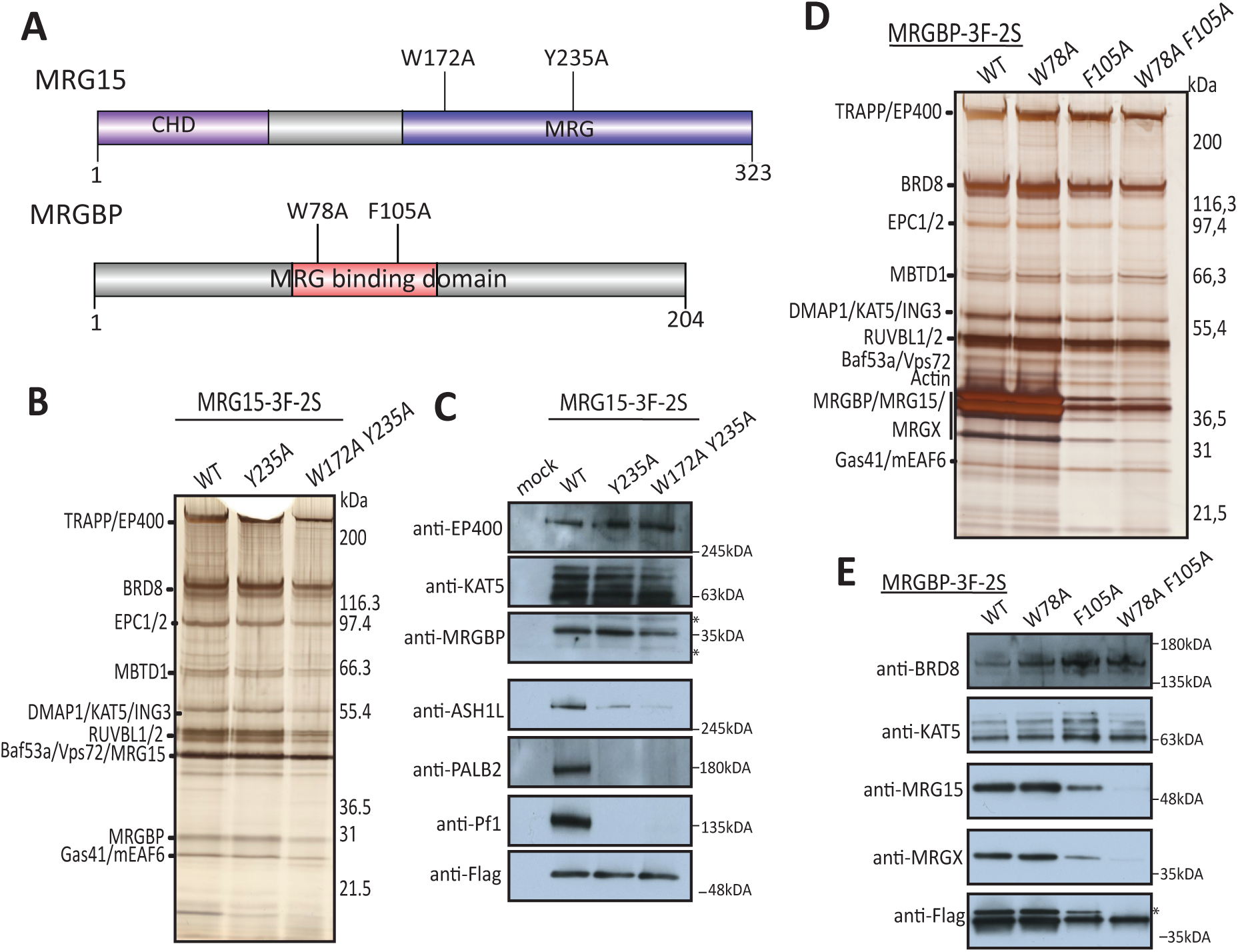
Structure-based mutations can differentially affect specific MRG15-containing complexes. (A) Schematic representation of the W172A Y235A mutations in MRG15 predicted to affect MRGBP/ASH1L/Pf1 interactions, and the W78A F105A mutations in MRGBP predicted to affect MRG15 interaction. (B) 3xFlag-2xStrep elutions of indicated MRG15 purifications from K562 cells were migrated on 4-12% SDS-PAGE and analyzed by silver staining. Strep elutions were analyzed by mass spectrometry in total spectrum count (Table S3). See also biological replicate in Figure S2B. (C) Purified complexes from (B) were analyzed by western blot with the indicated antibodies (* non-specific band). (D) 3xFlag-2xStrep elutions of indicated MRG15 purifications from K562 cells were migrated on 4-12% SDS-PAGE and analyzed by silver staining. (E) Purified complexes from (D) were analyzed by western blot with the indicated antibodies (* non-specific band).

### The MRG interactomes identify a new tetrameric protein complex

When we analyzed the interactomes of MRG15-S/L and MRGX, we noticed by gel, western and mass spectrometry the over-representation of specific subunits of NuA4/TIP60. (67)This was the case for MRGBP and the bromodomain-containing protein BRD8. This was confirmed by exponentially modified protein abundance index (emPAI)(67), similar to RUVBL1/2 subunits which are known to form a hexameric ring in protein complexes (**Fig. S3A-C**). That drew a parallel with a similar observation we made with yeast NuA4 components a few years ago (25). At that time, we noticed that MRG15 and MRGBP homologs, Eaf3 and Eaf7, were preferentially enriched when used as bait, along the protein Eaf5 which has no clear homolog in higher eukaryotes. We went on to characterize a yeast trimeric complex of Eaf5-Eaf7-Eaf3 that exists independently of NuA4, binds the coding region of active genes and interacts with the elongating RNAPII (25). TINTIN (<u>T</u>rimer Independent of <u>N</u>uA4 for <u>T</u>ranscription Interactions with <u>N</u>ucleosomes) is involved in transcription elongation presumably by facilitating disruption and recycling of nucleosomes from the front to the back of the elongating polymerase, also helping to suppress spurious transcription (68).

Since our purification of MRGBP, the human homolog of Eaf7, also yielded over overrepresentation of MRG15/X and BRD8 compared to the rest of NuA4/TIP60 subunits (**Fig. 3D, 4A**), we wondered if MRG15 in higher eukaryotes could also form an independent functional complex like yeast TINTIN, in which BRD8 would be the functional homolog of yeast Eaf5. To test this hypothesis, we first used Sf9 cells and co-infected them with baculovirus vectors coding for BRD8 (1BRD), MRG15 (S) and HA-tagged MRGBP. Fractionation of extracts with anti-HA beads and HA peptide elution clearly showed co-purification of BRD8 and MRG15 with MRGBP (**Fig. S2C**). We then generated K562 cell lines using the *AAVS1* system to purify BRD8 in comparison to MRGBP (**Fig. S4A**). Interestingly, BRD8 possesses two isoforms produced by alternative mRNA splicing, a shorter version with one H4ac-binding bromodomain (BRD) (Q9H0E9-2 to -4) and a longer version with two BRDs (Q9H0E9-1). Using the CRISPR/Cas9 system, we were able to tag both endogenous isoforms. We designed gRNAs to target the C-terminal of the two-BRD isoform or the N-terminal of the *BRD8* gene, tagging both isoforms (**Fig. S4B**). Although clonal selection and sequencing confirm the efficient introduction of our tag in the C-terminus of the long isoform, we were not able to clearly detect expression of this specific variant compared to the N-terminal tagging (**Fig. S4C,** maybe a very faint signal). Strangely, even expressing the 2BRD isoform from the *AAVS1* locus yielded very low signal, arguing that this isoform is unstable in K562 cells. Data in public databases confirm that expression of the 2BRD isoform is not really detected in most tissues except testis, while the 1BRD isoform is ubiquitously expressed (e.g. GTExportal.org).

**Figure 4.**
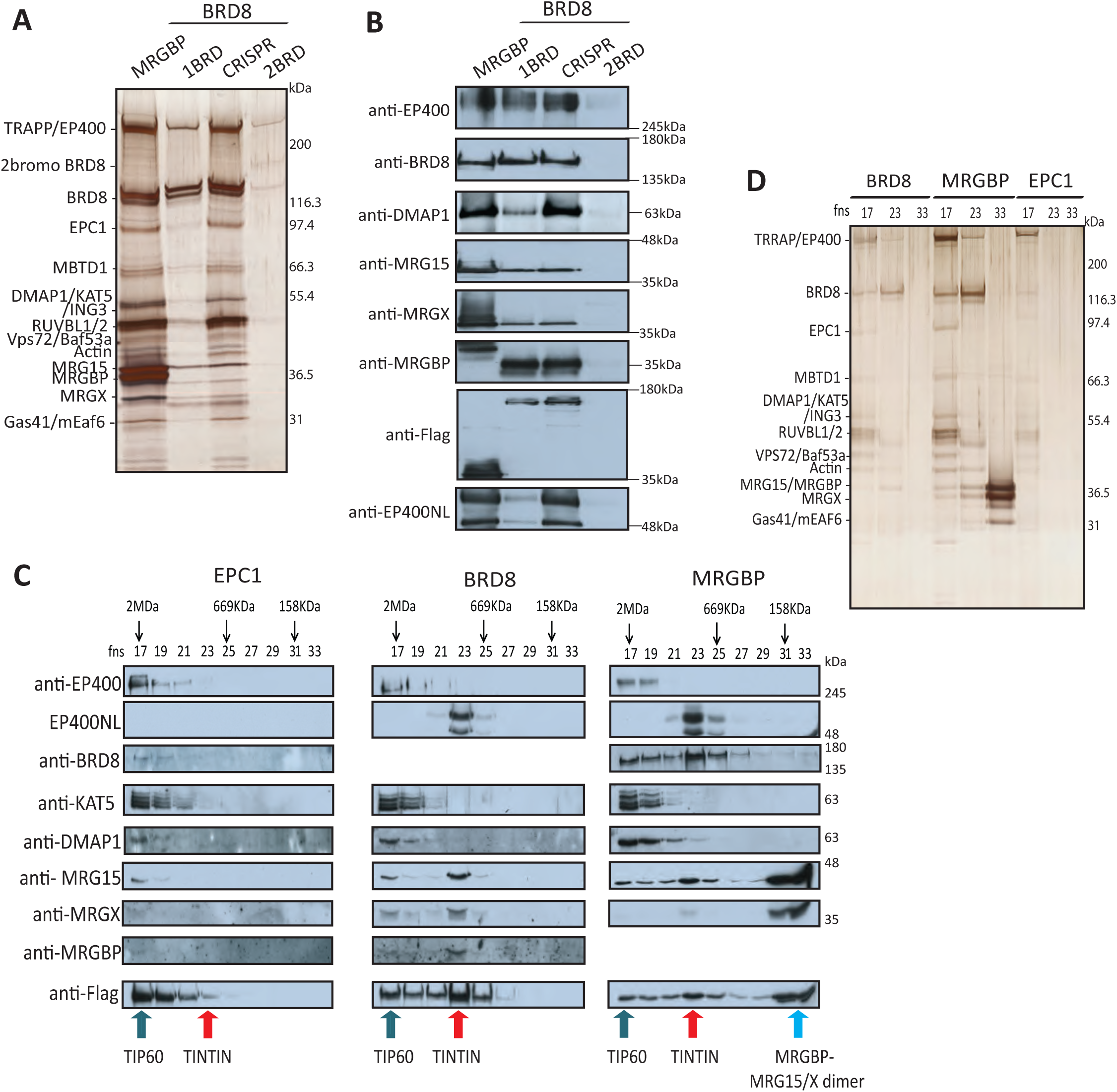
Identification of the TINTIN complex in human cells, composed of BRD8-MRGBP-MRG15/X and new subunit EP400NL. (A) Purification of native NuA4/TIP60 complex from K562 cells. Strep elutions of indicated purifications were analyzed on 4-12% SDS-PAGE followed by silver staining. MRGBP and spliced variants of BRD8 (1BRD, one bromodomain, 2BRD, two bromodomains, and CRISPR for N-term tagged BRD8) are linked to NuA4/TIP60 complex. (B) Western blots of purified complexes in (A) confirmed the association with the NuA4/TIP60 complex and the new subunit EP400NL. (C) Western blots of gel filtration fractions 17 to 33 (calibrated Superose 6, FPLC) with Flag elutions from EPC1, BRD8, and MRGBP *AAVS1* K562 cells (EPC1 purified fraction is used as a control reflecting the native NuA4/TIP60 complex (64)). Arrows point to specific annotated complexes with different elution profiles based on size. (D) 4-12% SDS-PAGE followed by silver staining of annotated fractions from (C) showing the NuA4/TIP60 complex (17), the TINTIN complex (23), and a dimer MRGBP/MRG15 or MRGX (33). See also Table S5 for mass spec results.

We tandem affinity purified MRGBP-3xFlag-2xStrep and BRD8-3xFlag-2xStrep expressed from *AAVS1* or endogenously tagged and analyzed the fractions by silver staining, mass spectrometry and western blots with the annotated antibodies (**Fig. 4A-B, Table S4**). Surprisingly, in those purifications, but also in our previous MRG15/X purifications, we detected an uncharacterized factor, called EP400 N-terminal like or EP400NL (**Fig. 1D and 4B**, **Tables S1, S3 and S4**). EP400NL was first described as a pseudogene and is homologous to a short region at the 5’ of the *EP400* gene (**Fig. S4D**). It can produce multiple spliced variants but only two of them (Q6ZTU2-5 and Q6ZTU2-6) with a specific C-terminal sequence were found associated with MRG proteins and BRD8 by mass spectrometry results (**Table S1, S3 and S4**). Thus, EP400NL proteins share a similar N-terminal sequence with the very large EP400 protein, an essential ATP-dependent remodeler and scaffolding subunit of the NuA4/TIP60 complex (**Fig. S4D**)(35, 69).

### EP400NL creates a human TINTIN complex by competing with EP400 for association to trimeric BRD8-MRGBP-MRG15/X

To prove the existence of this functional module independent of the NuA4/TIP60 complex, we purified BRD8 and MRGBP as above, but in parallel to the NuA4/TIP60-specific subunit EPC1 (64). We then loaded the tandem affinity-purified fractions on a Superose 6 size exclusion column to visualise distinct assemblies. Fractions from the gel filtration were analyzed by Western blot, silver-stained gel and mass spectrometry (**Fig. 4C-D**, **Table S5**). While the purified EPC1 fraction eluted early as expected with the large size of the NuA4/TIP60 (∼2MDa), BRD8 eluted as two populations with the strongest signal associated with a smaller protein complex (∼700–800 KDa) (**Fig. 4C-D**). This smaller population also co-eluted with the strongest signal of MRGBP and MRG15/X proteins. Importantly, in contrast to these proteins, EP400NL was only detected in the smaller size fraction, with no signal eluting with NuA4/TIP60. These results demonstrate the existence of a human TINTIN complex formed by EP400NL-BRD8-MRGBP-MRG15/X. Gel filtration of the purified MRGBP fraction gave similar results but with an important difference. In this case, a third much smaller population was detected below 150kDa and containing likely only MRGBP and MRG15/X (**Fig. 4C-D**, **Table S5**). Altogether, these data support the idea that the human TINTIN exists independently of the NuA4/TIP60 complex with potential functions in transcription as it was observed in yeast.

Since EP400NL was previously described as a pseudogene, we generated K562 cell lines using both *AAVS1* and CRISPR/Cas9 to, on one hand, tag the specific isoform Q6ZTU2-5 and, on the other hand, the endogenous gene at the C-terminus to tag the two specific isoforms found by mass spectrometry (**Fig. S4E-F**). We then used those cell lines to demonstrate physiological expression of the endogenous *EP400NL* gene and characterize the encoded protein interactome. Analysis of the tandem affinity-purified fractions confirmed expression of EP400NL by the endogenous gene and showed strong, apparently stoichiometric, association with BRD8, MRGBP and MRG15/X (**Fig. 5A-B, Table S6**)(note that MRGBP does not stain efficiently with silver, while showing strong signal by mass spectrometry)(see also emPAI analysis in **Fig. S3D**). Importantly, EP400NL did not purify with any other known components of NuA4/TIP60. Interestingly, an unusually high number of peptides from mRNA processing factors were found, suggesting a potential function of human TINTIN in mRNA processing or at least a close proximity during transcription elongation (**Table S6**).

**Figure 5.**
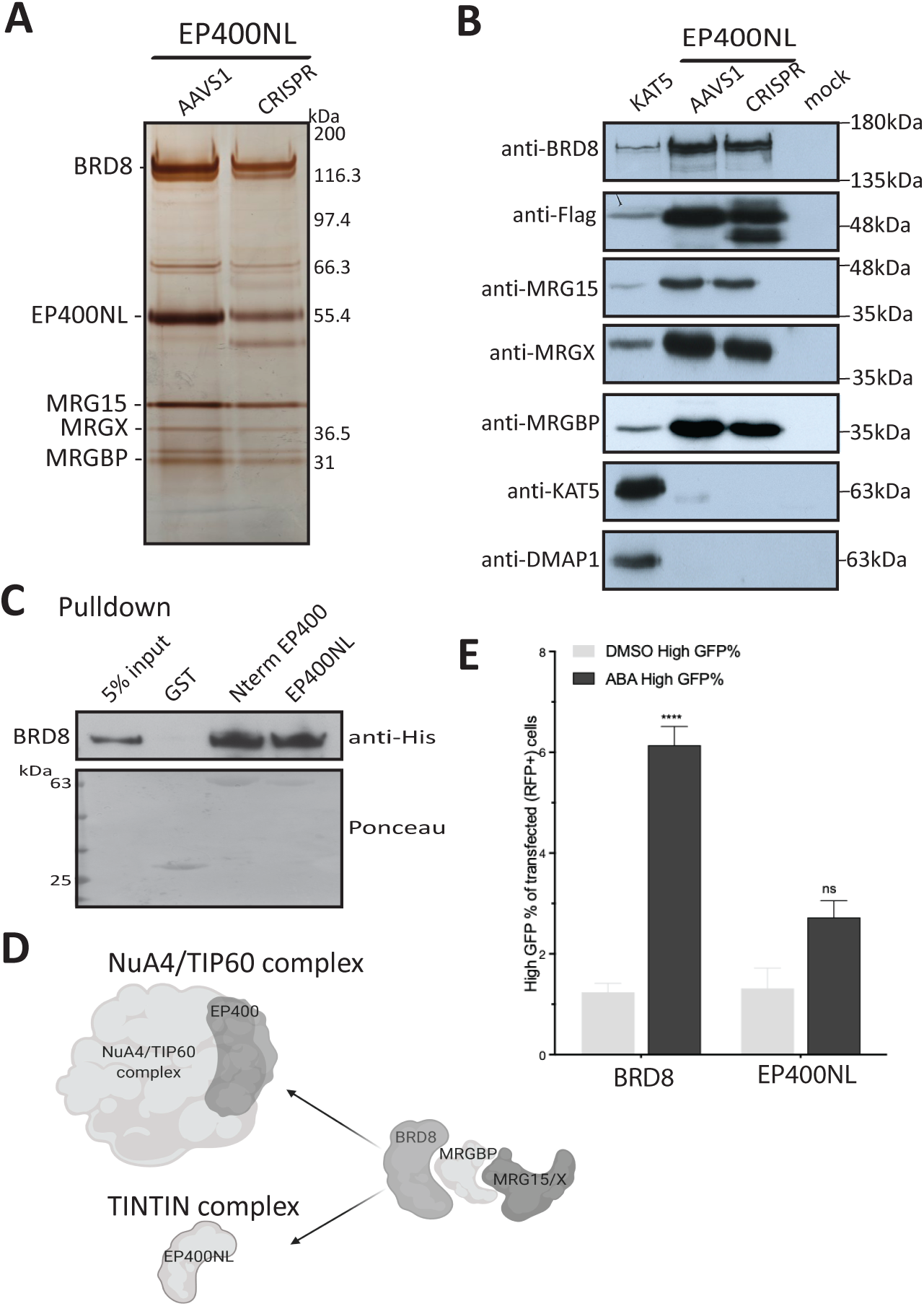
Endogenous EP400NL isolates TINTIN from NuA4/TIP60 through an interaction with BRD8. (A) 3xFlag-2xStrep elutions of Q6ZTU2-5 (*AAVS1*) or endogenous (CRISPR) EP400NL purified from K562 cells and analyzed on gel by silver staining. (B) Purified complexes from (A) were analyzed by western blot with the indicated antibodies to confirm the association of EP400NL with BRD8, MRGBP, and MRG15/X representing the human TINTIN complex. A KAT5/Tip60 purified fraction is used as a comparison for the NuA4/TIP60 complex (64). (C) BRD8 directly interacts with EP400NL and the homologous the N-terminal region of EP400 *in vitro*. Recombinant His-tagged BRD8 was incubated with the indicated GST fusion proteins on beads followed by washes and western blotting. (D) Schematic representation of the distinct associations of BRD8-MRGBP-MRG15/X with EP400 to form NuA4/TIP60 complex or EP400NL to form an independent TINTIN complex. (E) EP400NL is a weak transcriptional activator when artificially targeted to a promoter, in contrast to BRD8 which can be linked to the known co-activator function of NuA4/TIP60. Recruitment-activator reporter assay was quantified by flow cytometry analysis (47). It shows the % of cells transfected with EP400NL and BRD8 that express high level of GFP upon ABA-mediated targeting of the proteins to the reporter. At least 25,000 cells were analyzed for each replicate (mean ± s.e.m, n=3). DMSO is used as a negative control. Statistical analyses were performed by two-way ANOVA test followed by Tukey’s test, ****, p < 0.0001.

If EP400NL competes with EP400 to keep human TINTIN separated from the NuA4/TIP60 complex, it is important to identify its binding target. Direct physical interactions between BRD8 and MRGBP as well as MRGBP and MRG15 have been reported (20, 70). We have shown that Eaf5 is responsible for anchoring TINTIN to the yeast NuA4 complex, where it binds the first 85aa of Eaf1, the scaffolding subunit homologous to EP400 (25,71,72). Thus, if BRD8 is really the human functional homolog of yeast Eaf5, it should be responsible to bridge MRGBP-MRG15/X to EP400 in NuA4/TIP60 or EP400NL in TINTIN. To answer this question, we performed a pull-down assay with full-length recombinant BRD8 and GST fusions with homologous N-terminal regions of EP400 (aa194-446) and EP400NL (aa51-297) (see **Fig. S4D**). BRD8 was very efficiently pulled down by both fusions but not GST control, thereby confirming our hypothesis (**Fig. 5C**). Thus, BRD8 acts like the bridge that can anchors the MRG proteins to EP400 in the NuA4/TIP60 complex or to EP400NL to form the independent TINTIN complex. Thus, since the binding levels of BRD8 to EP400 N-terminus or EP400NL seem similar *in vitro* (**Fig. 5C**), relative protein abundance of EP400/EP400NL and BRD8 may be the determinants for the ratio of independent TINTIN complex in the cell (**Fig. 5D**). Analysis of public databases indicates that EP400NL mRNA is ubiquitously expressed, like EP400, but 3-fold less in average (e.g. GTExportal.org). To characterize a specific role of EP400NL/TINTIN in transcription, we used a recently described dCas9-based recruitment-activator reporter assay (47). This system measures transcription activation upon ABA-induced recruitment of factors at a promoter controlling the expression of GFP. Interestingly, EP400NL showed weak activator activity in this assay (**Fig. 5E**). In contrast, BRD8 showed significant transcription activation, likely in part due to the known co-activator function of NuA4/TIP60. This result suggests distinct functions of TINTIN and NuA4/TIP60 in the transcription process. As NuA4/TIP60 is known to function at gene promoters/start sites to favor transcription initiation (read-out of the assay in **Fig. 5E**), the weak effect of EP400NL may reflect a more important role on the body of genes to favor elongation, like we suggested for yeast TINTIN (25).

### The human TINTIN complex regulates both transcription and transcript isoforms of specific genes

The yeast TINTIN was shown to interact with the elongating RNAPII and to be involved in recycling disrupted nucleosomes in its wake to suppress spurious transcription (25). To test if human TINTIN also has a role in transcription, we performed siRNA-mediated knockdowns (KDs) of TINTIN components in comparison to KAT5/Tip60 knockdown, in an attempt to distinguish functions associated with NuA4/TIP60 vs independent TINTIN (**Fig. S5A-B**). RNAs were extracted 48h post-KDs and mRNA-sequencing was performed from samples depleted either for BRD8, MRGBP, MRG15 or KAT5/Tip60. 431 common genes were found downregulated in all KDs compared to siControl while 529 common genes were upregulated (**Fig. 6A**). These likely reflect the function of the NuA4/TIP60 complex. Due to the mutual exclusion of MRG15 and MRGX, we also compared KD of MRGX with the other KDs and found over 500 common genes downregulated or upregulated (**Fig. S5C**). Thus, some specific genes seem differently affected by the MRGX-containing NuA4/TIP60 complex versus the MRG15- containing one (1263 genes are commonly downregulated by MRG15/X KDs and 869 are upregulated, leaving two sets of nearly 2900 genes specifically affected by each one independently). Furthermore, when we excluded specific targets affected by the KD of KAT5/Tip60, we observed 161 downregulated genes and 124 upregulated genes commonly affected by KDs of TINTIN members BRD8, MRGBP and MRG15 (**Fig. 6A**). These numbers change to 253 downregulated and 142 upregulated when MRG15 KD is replaced by MRGX KD (**Fig. S5C**). Gene ontology term (GO-term) enrichment analyses were performed on those gene sets in order to identify specific biological processes and molecular function pathways in which TINTIN could be implicated (**Fig. 6B-C**, **S5D**). We also used RT-qPCR to validate some targets specifically regulated by the TINTIN complex but not the NuA4/TIP60 complex, such as *RFX8*, *HHIP*, *SYTL5*, *CD180*, which are downregulated by the KDs, and *AKT2* which is upregulated. *PRRG4* and *PCDHB9* are used as control where all KDs were similar to siControl (**Fig. 6D**). Interestingly, our ChIP-seq data (**Fig. S1C**), while in a different cell line, showed binding of MRG15/X to the *AKT2* gene.

**Figure 6.**
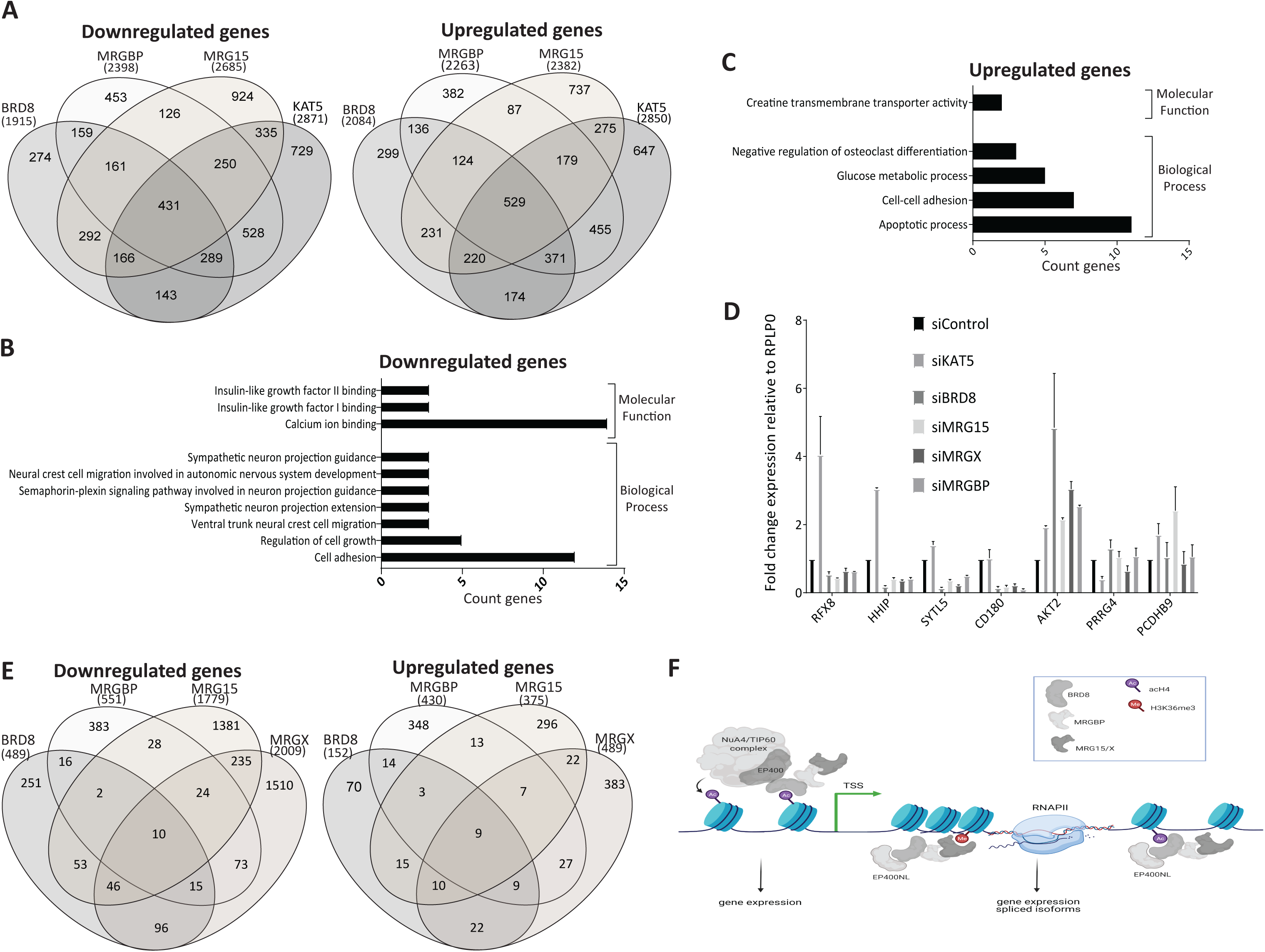
Human TINTIN is implicated in the expression of specific genes. (A) Venn diagrams showing that 161 genes are commonly downregulated (left) and 124 are upregulated in U2OS cells depleted (siRNA-mediated KDs) of BRD8, MRGBP and MRG15 but not in cells depleted of TIP60/KAT5. Genes showing high changes in expression, using a cutoff of 2-fold difference (Log2(fold change values), were tabulated between the selected KDs and control (siLuc). (B) and (C) Gene ontology analysis for downregulated (B) and upregulated genes (C) showing a significant enrichment (pValue <0.01) using DAVID 6.8. (D) RT-qPCR to validate target genes of TINTIN. *RFX8*, *HHIP*, *SYTL5*, and *CD180* are found downregulated, and *AKT2* is upregulated when TINTIN subunits are depleted. *PRRG4* and *PCDHB9* are used as control. Values represent means relative to control (siLuc). Error bars represent the range from two biological replicates. (E) Venn diagrams of downregulated (left) and upregulated (right) transcripts (versus genes) specifically regulated by BRD8, MRGBP, MRG15, or MRGX after exclusion of transcripts regulated by KAT5/TIP60, using a cutoff of 2-fold difference. siLuciferase was used as control. Two replicates of each KD were performed in U2OS cells. (F) Model of BRD8-MRGBP-MRG15/X function during transcription. BRD8 brings MRG proteins into the acetyltransferase NuA4/TIP60 complex due to its direct interaction with EP400 N-terminal region. This association is important for expression of specific gene targets. In addition, these subunits form a new independent functional module, called TINTIN, when BRD8 interacts instead with a new factor, EP400NL. TINTIN is also important for regulation of specific genes, maybe during the transcription elongation stage.

MRG15, its yeast homolog Eaf3 and MRGBP having already been linked to mRNA splicing processes (16-18,73), we determined if the ratio of alternative transcripts per gene were changed by KDs of TINTIN components. We first compared differential transcript expressions in KDs of BRD8, MRGBP, MRG15/X and KAT5/TIP60 to siControl. Then, we again excluded the transcripts that were significantly changed in KD KAT5/TIP60 (padj ≤ 0.1). The components of the TINTIN complex seem to commonly affect the regulation of only a few alternative transcripts (12 downregulated and 12 upregulated for TINTIN-MRG15, vs 25 down and 18 up for TINTIN-MRGX), even if each KD by itself shows a defect in the expression of much higher numbers of transcripts (**Fig. 6E**). Overall, these data demonstrate a significant function of the human TINTIN complex, independently of the NuA4/TIP60, in transcription regulation by modulating the expression of specific genes (**Fig. 6F**). We also showed that BRD8 and MRG proteins play also likely important roles in the regulation of some genes from within the NuA4/TIP60 acetyltransferase/remodeling complex or through colocalization on the same target of NuA4/TIP60 at the promoter and TINTIN on the coding region, as we have described in yeast (**Fig. 6F**)(25).

## Discussion

In this work, we characterized the interactomes of human MRG domain-containing proteins MRG15 and MRGX, factors that have been implicated in several distinct nuclear processes through association with diverse proteins. These experiments were done as to reflect the most native interactions, avoiding over-expression and when possible, using endogenous proteins as bait. We observed that MRG15 isoforms with long or short CHD and MRGX share the same interactome, such as stable association with the NuA4/TIP60 histone acetyltransferase complex and the Sin3B histone deacetylase/demethylase complex. Interestingly, MRG15 and MRGX are mutually exclusive in their associated complexes. Since MRGX does not possess the H3K36me3- binding chromodomain, it will be important to characterize the impact of this difference on the function of their partners. *MRG15* knock-out is embryonic lethal in mice, while *MRGX* knock-out does not lead to any obvious phenotype in development or cell proliferation, which may be in part due to some unidirectional compensation mechanism (19,74,75).

Interestingly, our data show that MRGX and MRG15 are stably associated with the endogenous H3K36me1/me2 methyltransferase ASH1L, in an apparent stoichiometric ratio since its activity cannot be stimulated *in vitro* by addition of exogenous MRG15 to relieve an auto-inhibitory loop (**Fig. 2**) (56–59). It will be interesting to study the functional impact of the MRG15 mutations that disrupt its association with endogenous ASH1L *in vivo*, through genome-wide location analysis of ASH1L and H3K36me2. The same mutants also disrupt association with the Sin3B complex. It was shown in yeast that Eaf3 plays a crucial role in the function of the homologous RPD3S complex to deacetylate nucleosomes in the wake of the elongating polymerase to suppress spurious transcription (24). A related function was proposed for the mammalian Sin3B complex (22, 76). Thus, it will be interesting to do transcriptomics to analyze if spurious transcription, i.e. cryptic initiation, anti-sense transcription, is detected with the MRG15 mutants. Since Sin3B also contains the KDM5 H3K4me3 demethylase that blocks the spreading of this mark from the TSS region to the body of the genes, it will also be important to verify the impact of the MRG15 mutant on the localization of this histone mark (76).

Most importantly, this study allowed the identification and characterization of a new functional MRG-containing complex, human TINTIN. We discovered a new partner of MRG15, EP400NL, previously dismissed as a probable pseudogene. EP400NL is key to create a BRD8-MRGBP-MRG15/X-containing TINTIN complex physically independent of NuA4/TIP60. EP400NL is homologous to the N-terminal region of NuA4/TIP60 scaffold subunit EP400, the interface on which the BRD8-MRGBP-MRG15/X trimer is normally anchored to the complex through BRD8. EP400NL can bind BRD8 presumably as efficiently as EP400 N-terminus and by doing so blocks the trimer from associating with NuA4/TIP60, creating an independent TINTIN complex. Interestingly, the mouse and human *EP400* genes are also known to produce a short splice variant encoding a truncated N-terminal protein corresponding to the same region where BRD8 binds, homologous to EP400NL (NCBI). This short splice variant of EP400 could have the same function as EP400NL to produce an NuA4/TIP60-independent TINTIN complex. Importantly, these proteins lack the HSA and SANT domains of EP400 that are required to assemble the NuA4/TIP60 complex (71, 72).

Since the yeast TINTIN complex is found on the coding region of active genes and plays a role during transcription elongation, we investigated if human TINTIN also affects gene expression. Indeed, several genes are deregulated in the absence of TINTIN components, independently of NuA4/TIP60 (**Fig. 6**). Further work will be necessary to better understand the function of TINTIN in mammals. EP400NL is likely an important target to use but, as mentioned above, EP400 short isoforms produced by alternative splicing may play a redundant role. Mapping the precise region of interaction between BRD8 and EP400NL/EP400 may be useful to design better tools to study TINTIN specific function.

Although MRG15 and MRGBP have been proposed to regulate mRNA splicing (16,17,73), we could not find a significant number of alternate transcripts specifically affected by the human TINTIN complex. Our mass spectrometry data also did not detect previously identified partners of MRG15 (PTB) and MRGBP (VEZF) for such function. It is possible that these interactions mostly occur on chromatin, making them difficult to detect in extracts. Independent KDs of MRG15, MRGX and MRGBP do show a larger number of transcripts being misregulated (**Fig. 6E**). This may be linked instead to the function of the small but abundant MRGBP-MRG15/X dimer that our gel filtration experiment identified (**Fig. 4**).

In conclusion, this study presented a highly confident native stable interactome of MRG proteins and uncovered a new mammalian complex implicated in gene regulation. It highlights the multi-specificity of the MRG domain for protein-protein interactions and associations with distinct protein complexes implicated in several chromatin-based nuclear processes. The presence of H4ac-binding bromodomain and H3K35me3-binding chromodomain in TINTIN also underlines again the cross-talk between transcription and chromatin modifications.

## Acknowledgments

We thank Suk Min Jang for work linked to this study and Tatiana Kutateladze for the recombinant ASH1L construct. This work was supported by grants from the Quebec Breast Cancer Foundation and the Canadian Institutes of Health Research (FDN-143314) to J.C., the Natural Sciences and Engineering Research Council of Canada (NSERC; 1304616-2017), the Cancer Research Society (25123) and Genome Quebec to J.P.L., University of Toronto startup funds to M.T. J.P.L. and S.I.H. were supported by Junior 1 salary awards from the Fonds de Recherche du Québec-Santé (FRQS) and J.C. held the Canada Research Chair in Chromatin Biology and Molecular Epigenetics.

## Data Availability

All MS files generated as part of this study were deposited at MassIVE (http://massive.ucsd.edu). The MassIVE ID is MSV000087245 and the MassIVE link for download is: http://massive.ucsd.edu/ProteoSAFe/status.jsp?task=9534fb8b77634860938b1139a53d9df8.

The password for download prior to final acceptance is MRG15.

ChIP-seq data from K562 cells and RNA-sequencing from U2OS cells were deposited in the GEO database under accession number GSE181533.

## Author contributions

MD, CR, CL, KJ and AP performed experiments. CJB and SH performed bioinformatic analyses. AL and JPL analyzed the proteomic data. AD, JPL and JC supervised and secured funding. MD and JC designed experiments, analyzed the data, and prepared the manuscript.

## Conflict of interest

The authors declare that they have no conflict of interest.

## Abbreviation

AP-MS: affinity purification combined to mass spectrometry
ASH1L: Absent, Small, or Homeotic discs 1-Like
BRD: bromodomain
CHD: chromodomain
CTD: C-terminal domain
EP400NL: EP400-N-terminal like
FPLC: Fast Protein Liquid Chromatography
K562: human erythroleukemic cells
KAT: lysine acetyltransferase
KDAC: lysine deacetylase
MRG15: MORF-related gene on chromosome 15
NuA4: Nucleosome Acetyltransferase of H4
RNAPII: RNA polymerase II
SETD2: SET Domain containing 2
TINTIN: Trimer Independent of NuA4 involved in Transcription Interactions with Nucleosomes
TIP60: Tat Interactive Protein 60kDa
U2OS: human osteosarcoma epithelial cells

---

**Table S1.**
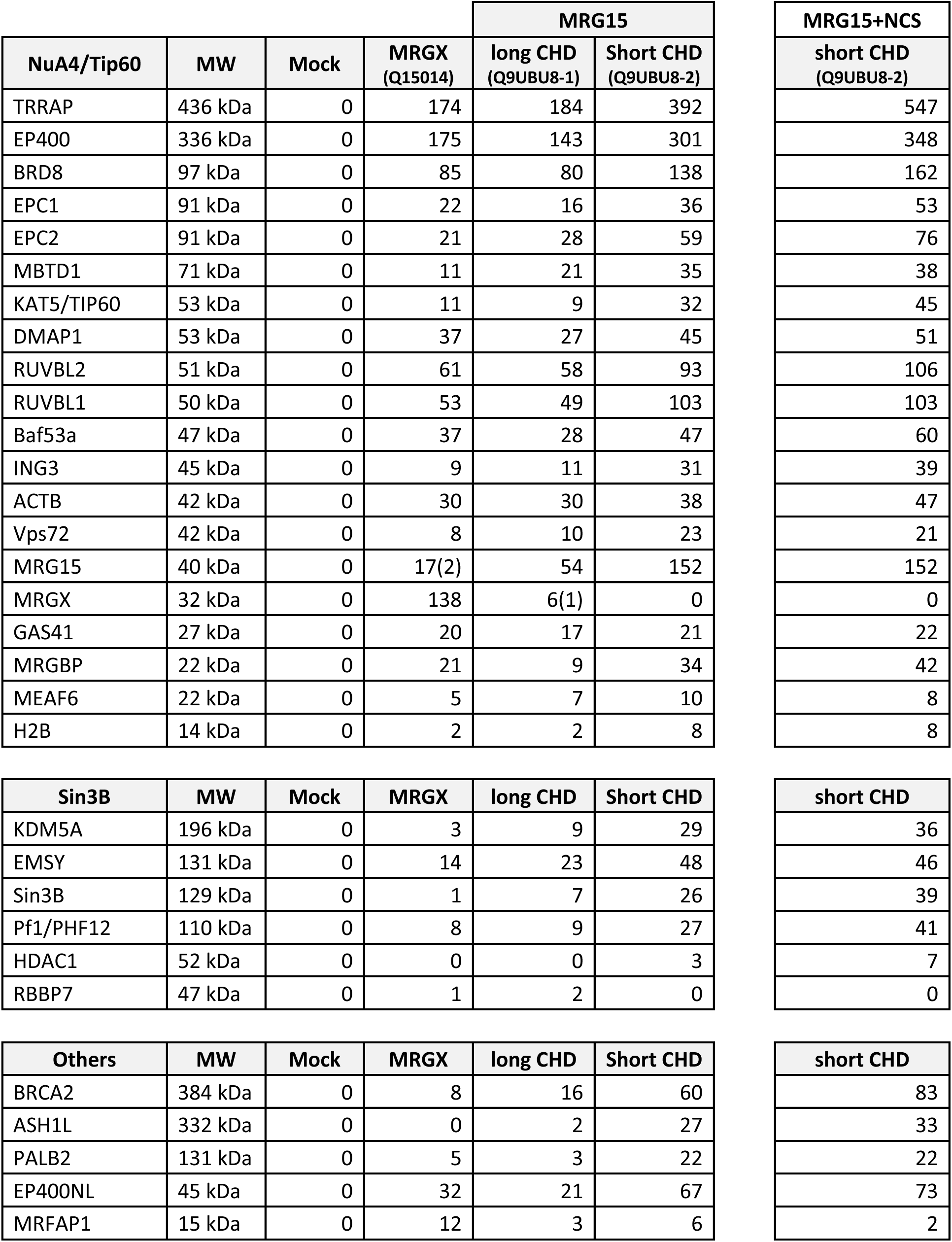
Mass spectrometry analysis of proteins that copurified with MRGX, MRG15 long and short CHD from K562 nuclear extracts and after DNA damage (NCS) (Total spectral counts), Related to Fig. 1, S2A. Numbers in parenthesis are the spectral counts specific (not shared peptide sequence) for MRG15 or MRGX found in each other fraction.

| NuA4/Tip60 | MW | Mock | MRGX<br>(Q15014) | MRG15 |  | MRG15+NCS<br>short CHD<br>(Q9UBU8-2) |
| --- | --- | --- | --- | --- | --- | --- |
|  |  |  |  | long CHD<br>(Q9UBU8-1) | Short CHD<br>(Q9UBU8-2) |  |
| TRRAP | 436 kDa | 0 | 174 | 184 | 392 | 547 |
| EP400 | 336 kDa | 0 | 175 | 143 | 301 | 348 |
| BRD8 | 97 kDa | 0 | 85 | 80 | 138 | 162 |
| EPC1 | 91 kDa | 0 | 22 | 16 | 36 | 53 |
| EPC2 | 91 kDa | 0 | 21 | 28 | 59 | 76 |
| MBTD1 | 71 kDa | 0 | 11 | 21 | 35 | 38 |
| KAT5/TIP60 | 53 kDa | 0 | 11 | 9 | 32 | 45 |
| DMAP1 | 53 kDa | 0 | 37 | 27 | 45 | 51 |
| RUVBL2 | 51 kDa | 0 | 61 | 58 | 93 | 106 |
| RUVBL1 | 50 kDa | 0 | 53 | 49 | 103 | 103 |
| Baf53a | 47 kDa | 0 | 37 | 28 | 47 | 60 |
| ING3 | 45 kDa | 0 | 9 | 11 | 31 | 39 |
| ACTB | 42 kDa | 0 | 30 | 30 | 38 | 47 |
| Vps72 | 42 kDa | 0 | 8 | 10 | 23 | 21 |
| MRG15 | 40 kDa | 0 | 17(2) | 54 | 152 | 152 |
| MRGX | 32 kDa | 0 | 138 | 6(1) | 0 | 0 |
| GAS41 | 27 kDa | 0 | 20 | 17 | 21 | 22 |
| MRGBP | 22 kDa | 0 | 21 | 9 | 34 | 42 |
| MEAF6 | 22 kDa | 0 | 5 | 7 | 10 | 8 |
| H2B | 14 kDa | 0 | 2 | 2 | 8 | 8 |

| Sin3B | MW | Mock | MRGX | long CHD | Short CHD | short CHD |
| --- | --- | --- | --- | --- | --- | --- |
| KDM5A | 196 kDa | 0 | 3 | 9 | 29 | 36 |
| EMSY | 131 kDa | 0 | 14 | 23 | 48 | 46 |
| Sin3B | 129 kDa | 0 | 1 | 7 | 26 | 39 |
| Pf1/PHF12 | 110 kDa | 0 | 8 | 9 | 27 | 41 |
| HDAC1 | 52 kDa | 0 | 0 | 0 | 3 | 7 |
| RBBP7 | 47 kDa | 0 | 1 | 2 | 0 | 0 |

| Others | MW | Mock | MRGX | long CHD | Short CHD | short CHD |
| --- | --- | --- | --- | --- | --- | --- |
| BRCA2 | 384 kDa | 0 | 8 | 16 | 60 | 83 |
| ASH1L | 332 kDa | 0 | 0 | 2 | 27 | 33 |
| PALB2 | 131 kDa | 0 | 5 | 3 | 22 | 22 |
| EP400NL | 45 kDa | 0 | 32 | 21 | 67 | 73 |
| MRFAP1 | 15 kDa | 0 | 12 | 3 | 6 | 2 |

**Table S2.**
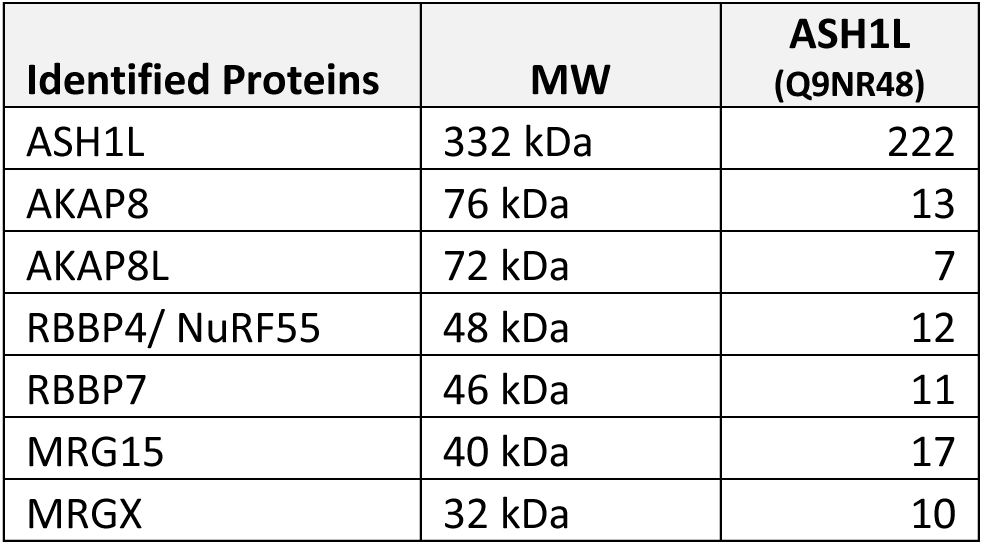
Mass spectrometry analysis of endogenous ASH1L purified from K562 cells (total spectral counts). Related to Figure 2.

| **Identified Proteins** | **MW** | **ASH1L**<br>**(Q9NR48)** |
| --- | --- | --- |
| ASH1L | 332 kDa | 222 |
| AKAP8 | 76 kDa | 13 |
| AKAP8L | 72 kDa | 7 |
| RBBP4/ NuRF55 | 48 kDa | 12 |
| RBBP7 | 46 kDa | 11 |
| MRG15 | 40 kDa | 17 |
| MRGX | 32 kDa | 10 |

**Table S3.** Mass spectrometry analysis of MRG15 WT versus mutant interactomes purified from K562 cells through MRG15-3xFlag 2xStrep integrated at AAVS1 (total spectral counts). Related to Figures 3, S2B.

|  | Identified proteins | MW | MRG15 |  |  |  |  |
| --- | --- | --- | --- | --- | --- | --- | --- |
|  |  |  | WT #1 | WT #2 | Y235A | W172A Y235A #1 | W172A Y235A #2 |
| **NuA4/TIP60 complex** | TRRAP | 436 kDa | 143 | 703 | 43 | 22 | 495 |
|  | EP400 | 343 kDa | 240 | 462 | 210 | 61 | 349 |
|  | BRD8 | 103 kDa | 105 | 231 | 86 | 35 | 173 |
|  | EPC2 | 91 kDa | 51 | 137 | 34 | 14 | 82 |
|  | EPC1 | 91 kDa | 29 | 71 | 23 | 7 | 49 |
|  | MBTD1 | 71 kDa | 15 | 57 | 6 | 1 | 29 |
|  | KAT5 | 62 kDa | 28 | 99 | 13 | 5 | 75 |
|  | DMAP1 | 53 kDa | 52 | 92 | 17 | 14 | 69 |
|  | RUVBL2 | 51 kDa | 166 | 203 | 79 | 34 | 108 |
|  | RUVBL1 | 50 kDa | 90 | 199 | 62 | 18 | 151 |
|  | ACTL6A | 47 kDa | 45 | 66 | 30 | 16 | 48 |
|  | ING3 | 45 kDa | 19 | 57 | 3 | 2 | 33 |
|  | ACTB | 42 kDa | 45 | 58 | 25 | 64 | 50 |
|  | VPS72 | 41 kDa | 26 | 36 | 6 | 7 | 28 |
|  | MRG15 | 37 kDa | 203 | 173 | 163 | 89 | 178 |
|  | YEATS4 | 27 kDa | 15 | 33 | 4 | 4 | 20 |
|  | MRGBP | 22 kDa | 27 | 55 | 27 | 9 | 28 |
|  | MEAF6 | 22 kDa | 6 | 22 | 0 | 0 | 15 |
| **TINTIN** | EP400NL | 52 kDa | 62 | 72 | 39 | 14 | 59 |
| **Sin3B complex** | EMSY | 141 kDa | 48 | 50 | 0 | 0 | 0 |
|  | PHF12/Pf1 | 110 kDa | 22 | 42 | 0 | 0 | 0 |
|  | SIN3B | 133 kDa | 9 | 39 | 0 | 0 | 0 |
|  | KDM5A | 192 kDa | 9 | 54 | 0 | 0 | 0 |
|  | RBBP4 | 48 kDa | 4 | 7 | 0 | 0 | 1 |
| **Other factors** | BRCA2 | 384 kDa | 21 | 71 | 0 | 0 | 0 |
|  | ASH1L | 333 kDa | 10 | 26 | 0 | 0 | 1 |
|  | PALB2 | 131 kDa | 14 | 31 | 0 | 0 | 0 |
|  | MRFAP1 | 15 kDa | 6 | 0 | 0 | 0 | 0 |

**Table S4.** Mass spectrometry analysis of NuA4/TIP60 complex and EP400NL that interact with MRGBP and BRD8 transcripts (with 2BRDs or all transcripts (N-term CRISPR) or 1BRD), purified from K562 cells (total spectral counts). Related to Figure 4.

| Identified Proteins | MW | BRD8 |  |  | MRGBP<br>(Q9NV56) |
| --- | --- | --- | --- | --- | --- |
|  |  | Endogenous | 1 BRD<br>(Q9H0E9-2) | 2 BRD<br>(Q9H0E9-2) |  |
| TRRAP | 436 kDa | 330 | 110 | 33 | 408 |
| EP400 | 340 kDa | 275 | 139 | 42 | 340 |
| BRD8 | 135 kDa | 117 | 108 | 23 | 168 |
| EPC2 | 91 kDa | 68 | 31 | 5 | 63 |
| EPC1 | 91 kDa | 33 | 18 | 2 | 45 |
| MBTD1 | 71 kDa | 30 | 16 | 3 | 34 |
| KAT5 | 53 kDa | 20 | 13 | 0 | 29 |
| DMAP1 | 53 kDa | 59 | 32 | 7 | 60 |
| RUVBL2 | 51 kDa | 127 | 62 | 26 | 137 |
| RUVBL1 | 50 kDa | 84 | 35 | 18 | 118 |
| ACTL6A | 47 kDa | 47 | 23 | 8 | 51 |
| ING3 | 45 kDa | 23 | 4 | 0 | 34 |
| ATCB | 42 kDa | 46 | 28 | 11 | 49 |
| VPS72 | 42 kDa | 14 | 12 | 3 | 22 |
| MRG15 | 40 kDa | 44 | 54 | 7 | 251 |
| MRGX | 32 kDa | 42 | 56 | 9 | 142 |
| YEATS4 | 27 kDa | 27 | 15 | 3 | 25 |
| MRGBP | 22 kDa | 23 | 25 | 5 | 115 |
| MEAF6 | 22 kDa | 8 | 8 | 0 | 9 |
| H2B | 14 kDa | 0 | 7 | 0 | 4 |
| EP400NL | 45 kDa | 64 | 11 | 0 | 70 |

**Table S5.** Mass spectrometry analysis of FPLC gel filtration fractions 23 of BRD8 and MRGBP corresponding to the TINTIN complex and 33 of MRGBP corresponding to the MRGBP/MRG15 or MRGX dimer (total spectral counts). Related to Figure 4.

| Identified Proteins | MW | BRD8 | MRGBP |  |
| --- | --- | --- | --- | --- |
|  |  | Fraction 23 | Fraction 23 | Fraction 33 |
| BRD8 | 103 kDa | 76 | 110 | 8 |
| MRG15 | 41 kDa | 9 | 39 | 15 |
| EP400NL | 38 kDa | 23 | 56 | 3 |
| MRGX | 32 kDa | 12 | 35 | 20 |
| MRGBP | 22 kDa | 16 | 30 | 24 |

**Table S6.** Mass spectrometry analysis of EP400NL interactome purified from K562 cells through EP400NL-3xFlag-2xStrep integrated at AAVS1 or from CRISPR/Cas9-mediated tagging of the endogenous gene (total spectral counts). Related to Figure 5.

| TINTIN | MW | EP400NL |  |
| --- | --- | --- | --- |
|  |  | CRISPR | Q6ZTU2-5 AAVS1 |
| BRD8 | 103 kDa | 192 | 313 |
| EP400NL | 52 kDa | 95 | 133 |
| MRG15 | 37 kDa | 117 | 164 |
| MRGX | 32 kDa | 71 | 107 |
| MRGBP | 22 kDa | 41 | 63 |

| mRNA processing factors | MW | EP400NL |  |
| --- | --- | --- | --- |
|  |  | CRISPR | Q6ZTU2-5 AAVS1 |
| SRRM2 | 300 kDa | 11 | 4 |
| SF3B1 | 146 kDa | 5 | 2 |
| SF3B3 | 136 kDa | 13 | 13 |
| GEMIN4 | 119 kDa | 2 | 0 |
| U2SURP | 118 kDa | 6 | 6 |
| SF3B2 | 100 kDa | 7 | 8 |
| RBM5 | 92 kDa | 5 | 2 |
| DDX20 | 92 kDa | 2 | 1 |
| HNRNPM | 78 kDa | 14 | 15 |
| PRPF31 | 55 kDa | 12 | 6 |
| HNRNPK | 51 kDa | 7 | 4 |
| HNRNPF | 46 kDa | 3 | 5 |
| RBM17 | 45 kDa | 2 | 4 |
| HNRNPC | 32 kDa | 2 | 0 |
| SNRPD3 | 14 kDa | 2 | 1 |

Note: All protein identifications determined by mass spec of different fractions are included in supplementary tables within a **supplemental Excel file**, including accession numbers, spectral counts, unique peptides and percentage coverage.

## SUPPLEMENTAL EXPERIMENTAL PROCEDURES

### ChIP-sequencing Experiments

For Flag ChIP, 1mg of cross-linked chromatin from K562 cells was incubated with 10µg of anti-Flag antibody (Sigma, M2) pre-bound on 300 µl of Dynabeads Prot-G (Invitrogen) overnight at 4°C. The beads were washed extensively and eluted in 0.1% SDS, 0.1M NaHCO3. Crosslink was reversed with 0.2M NaCl and incubation overnight at 65C. Samples were treated with RNase and Proteinase K for 2h and recovered by phenol-chloroform and ethanol precipitation. Libraries for sequencing were prepared with TruSeq LT adaptors (Illumina). Samples were sequenced by 50 bp single reads on HiSeq 4000 platform (Illumina).

### Processing, Alignment and Peak Calling of ChIP-seq Data

FastQ format reads were aligned to the hg19 human reference using the Bowtie alignment algorithm (1). Bowtie2 version 2.1.6 was used with the pre-set sensitive parameter to align ChIP sequencing reads. MACS version 2.0.10 (model-based analysis of ChIPseq) peak finding algorithm was used to identify regions of ChIP-Seq enrichment over the background (2). The pipeline, commands, and parameters that were used are: Trimming of sequence (filter out 39 adaptor, and remove last 2 bases and 3 extra bases if it matches with adaptor sequence). Mapping sequences to human genome (hg19) using Bowtie: (i) command: bowtie2 -p 8 –sensitive -x genome/genome -U sequence.reads.fastq – S sample.sam. Peak calling algorithm MACS: (i) command: macs2 callpeak –t ChIPseq.data.bam -c input.sample.file.bam -- broad -f BAM -g hs -n [directory] –outdir MACS2_files --nomodel --shiftsize 100 –B. Unique mapped read values were normalized to library size. Peaks were annotated as per human genome EnsDb.Hsapiens.v75 – hg19. Raw sequences and processed data of ChIP-sequencing from K562 cells were deposited in the GEO database under accession number **GSE181533**.

### Cell Cycle Analysis

Fluorescence-activated cell sorting (FACS) analysis was used for cell cycle profiling. The cells were harvested by trypsinization, fixed with 70% ethanol, treated with propidium iodide for FACS analysis on a BD Accuri C6 Plus.

### Baculovirus expression

BRD8, MRG15 and MRGBP cDNA were cloned in pFastBac vectors, packaged in viral particles with DH10Bac competent cells, and used to co-infected Sf9 cells. 48hrs post-infection cells were harvested, protein extracted and incubated with anti-HA beads. After washes, HA peptides were used to elute HA-tagged MRGBP and its associated proteins.

## SUPPLEMENTAL FIGURE LEGENDS

**Figure S1.**
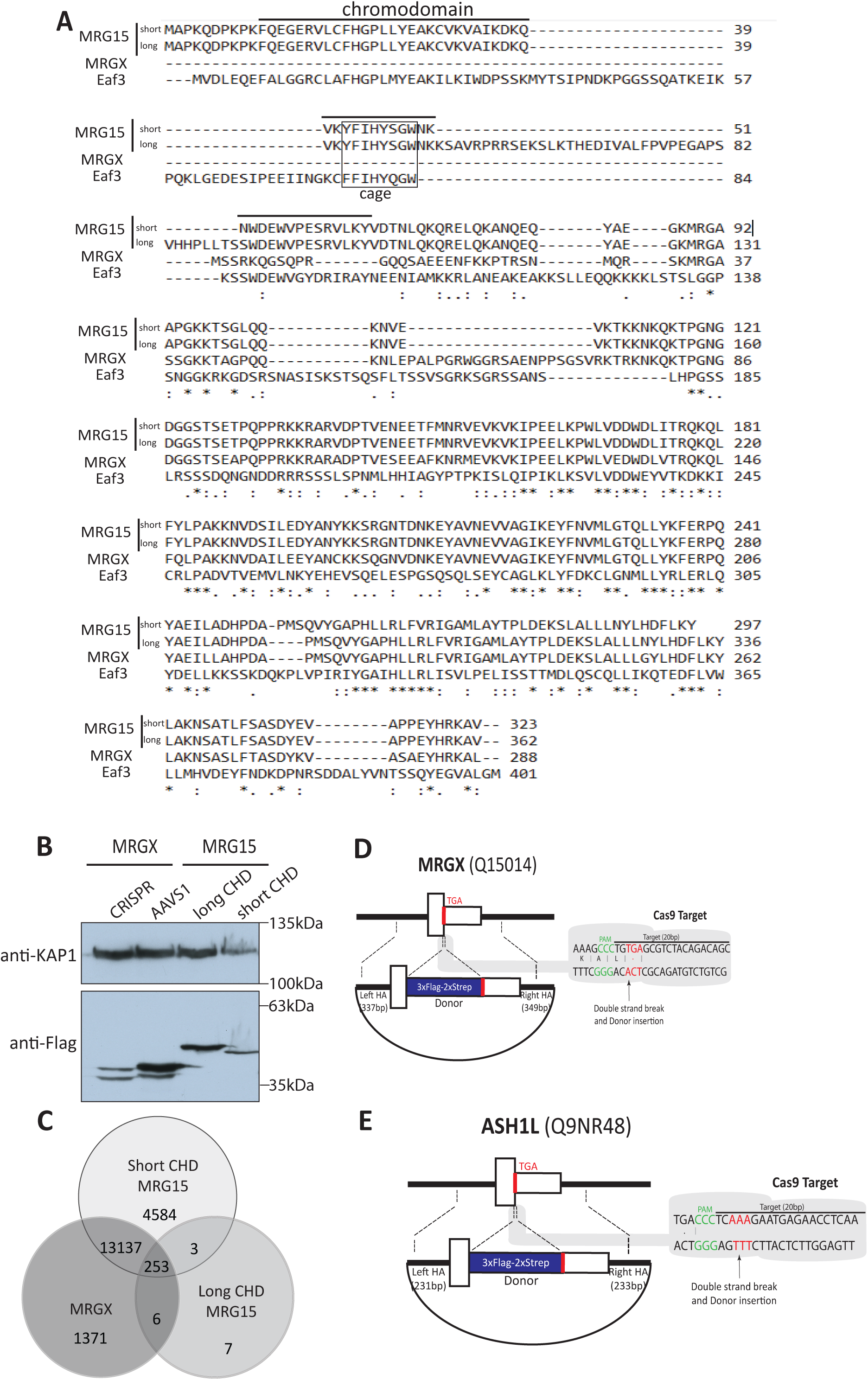
MRG15 isoforms and MRGX binding to chromatin *in vitro* and *in vivo*, related to Figures 1 and 2. (A) Amino acids sequence alignment of MRG15 isoforms that have a “long” or “short” chromodomain, the paralog MRGX without chromodomain, and the Eaf3 homolog from *Saccharomyces cerevisiae*. The chromodomain and hydrophobic cage necessary to bind methylated lysine 36 of histone H3 are indicated. (B) Whole-cell extract of isogenic K562 cells stably expressing the indicated proteins with a 3xFlag-2xStrep tag from the *AAVS1* safe harbor locus or endogenously tagged (CRISPR MRGX). (C) Anti-FLAG ChIP-seq analysis of MRG15-S/L and MRGX in K562 cells using the *AAVS1* isogenic cell lines. Venn diagram of mapped significant genes that are bound for each protein and overlaps. MRG15 short isoform (MRG15-S) and MRGX bind mostly the same loci. Only a very small number of bound loci was detected for the MRG15 long isoform. Numbers represent the number of significant genes in the vicinity of mapped peaks. (D) Genomic editing with the CRISPR/Cas9 system for tagging the endogenous MRGX. Schematic of the *MRGX* locus, Cas9 target site, and donor construct used to insert the cassette containing 3xFlag-2xStrep in the last exon. Annotated are the positions of the stop codon target site, the PAM motif, and homology arms left and right (HAL and R). (E) Schematic of the *ASH1L* locus, Cas9 target site, and donor construct used to insert the cassette containing 3xFlag-2xStrep in the last exon. Annotated are the positions of the stop codon target site, the PAM motif, and homology arms left and right (HAL and R).

**Figure S2.**
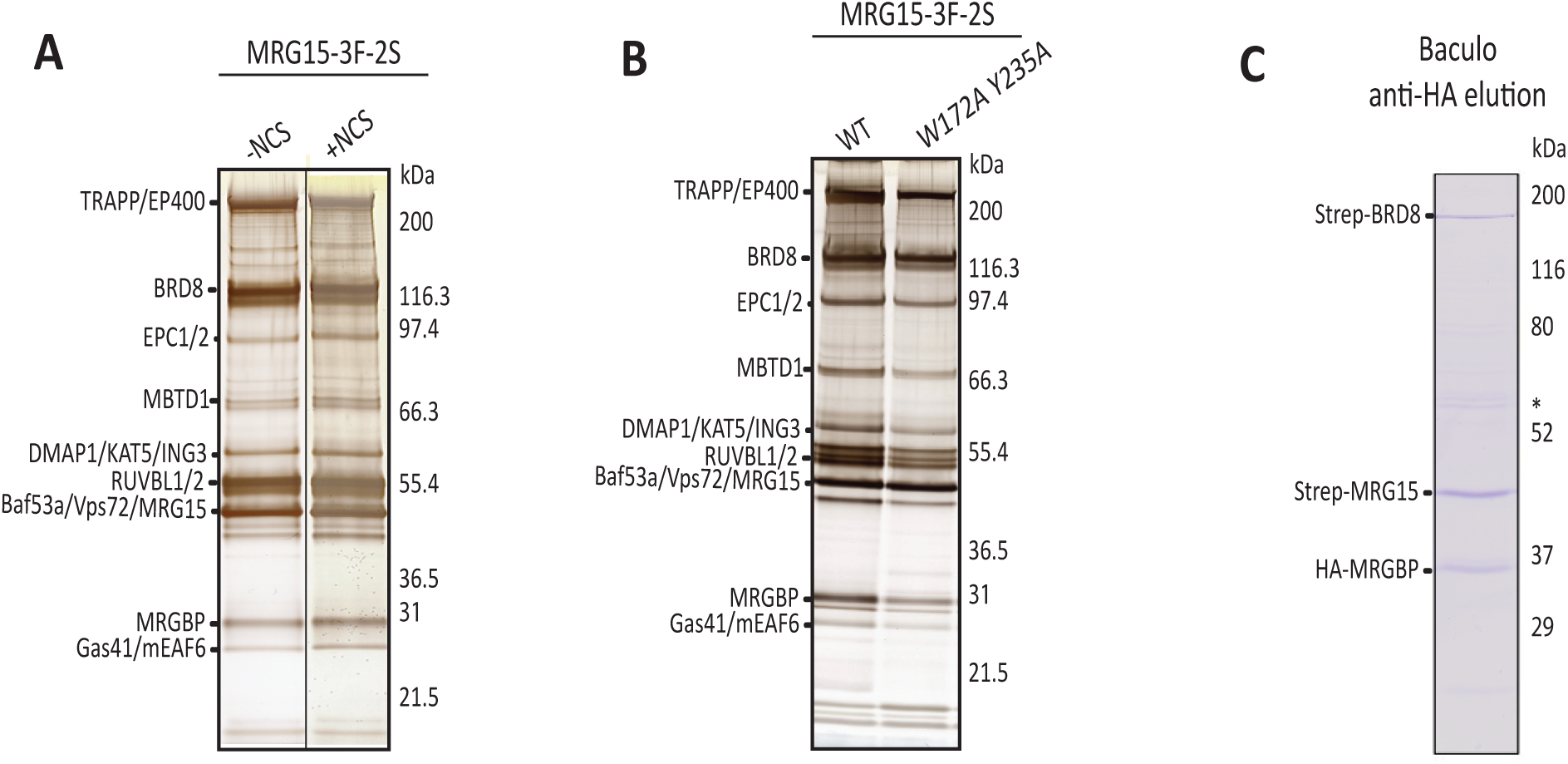
MRG15 purification after DNA damage or carrying mutations and baculovirus expression of the BRD8-MRGBP-MRG15 trimer, related to Figures 1, 3, 4 and Tables S1, S3. (A) Tandem affinity purified MRG15 (short) from K562 cells treated, or not, with 50ng/mL of neocarzinostatin (NCS) for 3h before being collected to prepare the nuclear extracts. Fractions were run on gel and silver stained. (B) Tandem affinity purified wild-type (WT) or double mutant (W172A Y235A) MRG15 (short) from K562 cells. Fractions were run on gel and silver stained. (C) Purification of reconstituted BRD8-MRGBP-MRG15 trimer from baculovirus. Sf9 cells were coinfected with single viruses for 48h. Single HA immunopurification was followed by elution with HA peptides. Purified fraction was migrated on gel and stained with coomassie. (*contaminant).

**Figure S3.**
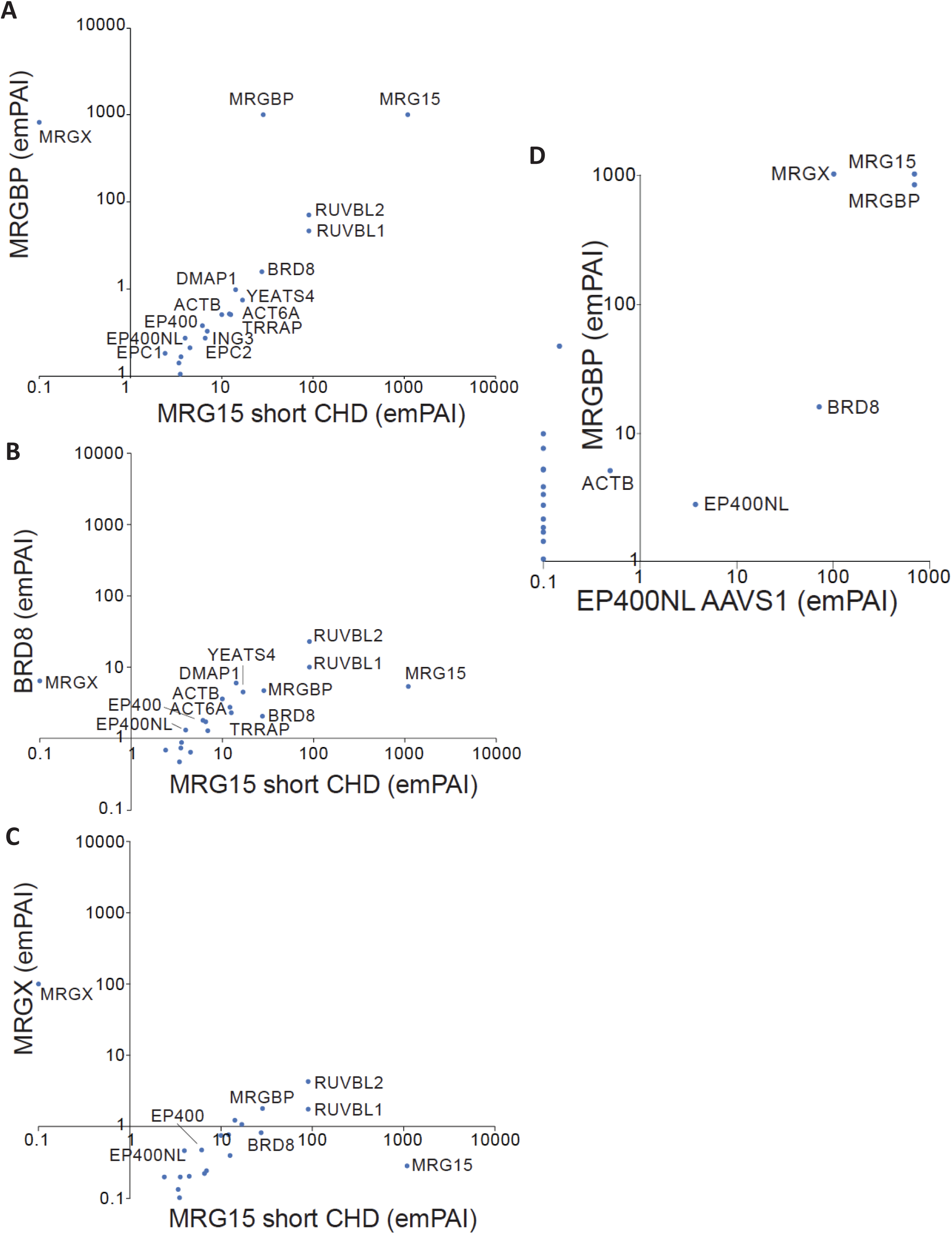
Exponentially modified protein abundance index (emPAI) of MRG15, MRGBP, BRD8, MRGX and EP400NL purified fractions to evaluate ratio of interactors, related to Figures 1, 3, 4 and 5, Tables S1, S3, S4 and S6. (A-C) emPAI of MRGBP vs MRG15, BRD8 vs MRG15 and MRGX vs MRG15. The higher abundance of MRGBP, BRD8 with MRG15/X supports the existence of a trimeric complex. RUVBL1/2 are also abundant since they form a hexameric ring within NuA4/TIP60. (D) emPAI of MRGBP vs EP400NL to support the concept of a stoichiometric assembly of EP400NL-BRD8-MRGBP-MRG15/X.

**Figure S4.**
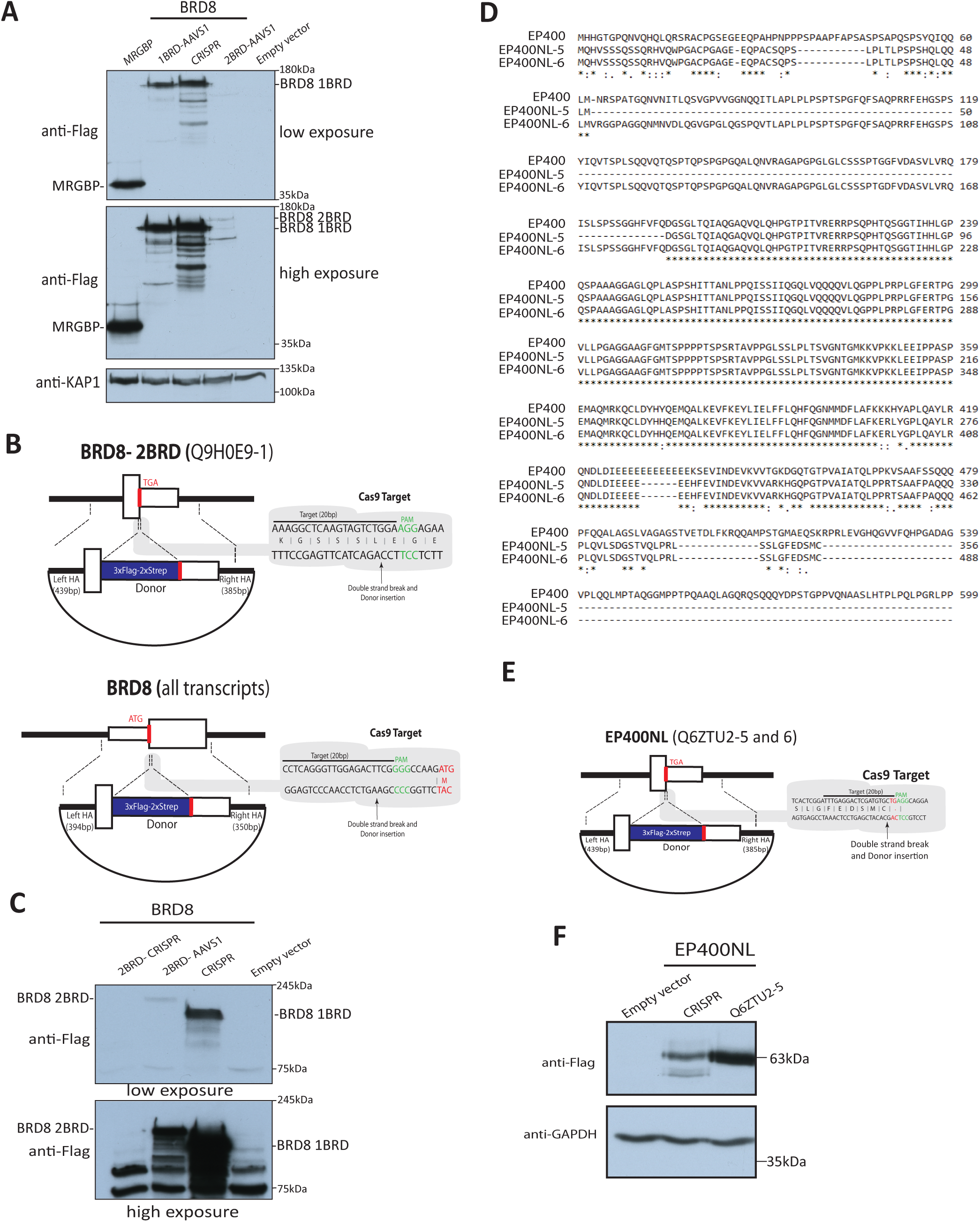
CRISPR/Cas9-mediated tagging of endogenous BRD8 and EP400NL, related to Figures 4-5. (A) Whole-cell extracts (WCE) were analyzed by immunoblotting with anti-Flag. Anti-KAP1 was used as loading control. (B) Genomic editing with the CRISPR/Cas9 system for tagging the double bromodomain (2BRD) isoform BRD8 (at the C-term) and all BRD8 isforms (at the N-term). Schematic of the *BRD8* locus, Cas9 target site, and donor construct used to insert the cassette containing 3xFlag-2xStrep in the last exon for 2BRD or first exon for all isoforms. Annotated are the positions of the stop/start codon target sites, the PAM motif, and homology arms left and right (HAL and R). (C) Anti-Flag analysis of BRD8 whole-cell extracts from *AAVS1* and endogenously tagged cell lines. The N-terminal endogenous tagging reveals the main expression of the 1BRD isoform, and the C-terminal tagging fails to detect a clear signal. Even expression from AAVS1 leads to low level of the 2BRD isoform. (D) Protein sequence alignment to show the very high similarity/identity of the N-terminal region of human EP400 with the previously uncharacterized EP400NL protein. (E) Genomic editing with the CRISPR/Cas9 system for tagging endogenous EP400NL in C-term. Schematic of the *EP400NL* locus, Cas9 target site, and donor construct used to insert the cassette containing 3xFlag-2xStrep in the last exon. Annotated are the positions of the stop codon target site, the PAM motif, and homology arms left and right (HAL and R). (F) EP400NL expression from endogenously tagged or *AAVS1* K562 cells. Anti-GAPDH is used as loading control.

**Figure S5.**
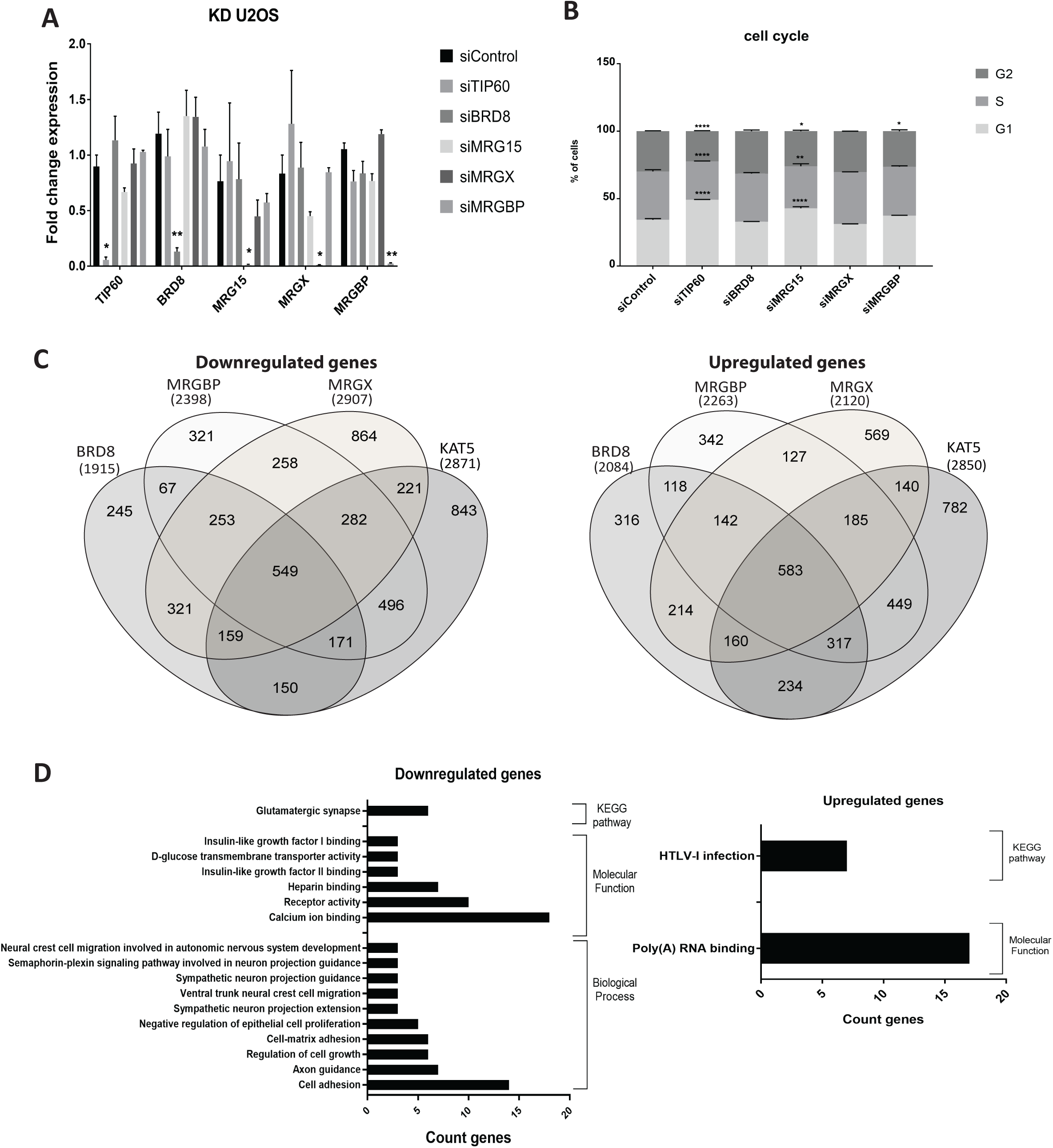
Depletion of TIP60 versus TINTIN subunits in U2OS cells, related to Figure 6. (A) Validation by RT-qPCR of the different siRNAs used for depletion and RNA sequencing analyses. RPLP0 was used as control. Different sets of primers were annotated. Knockdowns (KDs) were normalized with the siControl (siLuciferase). Error bars represent the range of two independent experiments. (B) Cell cycle profile analysis of U2OS cells 48hrs after transfection of the different siRNAs. Error bars represented the range of two independent experiments. Statistical analyses were performed by two-way ANOVA test followed by Tukey’s test, *, p < 0.1, **, p < 0.01, ****, p < 0.0001. (C) 253 genes are downregulated (left) and 142 genes are upregulated (right) by KDs of TINTIN subunits (BRD8, MRGBP, and MRGX) but not NuA4/TIP60 complex. To highlight the genes with high changes in expression, a cutoff of 2-fold difference was applied to the Log2(fold change) values between the selected KDs and control (siLuc). (D) Gene ontology analysis for downregulated (left) and upregulated genes (right) showing a significant enrichment (pValue <0.01) using DAVID 6.8.

## Notes

### Competing Interest Statement

The authors have declared no competing interest.

